# Solid-like PAR protein assemblies encode long-term spatial memory of cell polarity during suspended animation

**DOI:** 10.64898/2026.06.30.733387

**Authors:** Joana Borrego-Pinto, Jake Cornwall Scoones, Nisha Hirani, KangBo Ng, Buzz Baum, Shiladitya Banerjee, Nathan W. Goehring

## Abstract

During suspended animation, organisms must preserve cellular organization despite the collapse of the active biochemical processes that normally maintain it. Here we show that anoxia-induced suspended animation drives cell polarity proteins into a poised, memory-like state that preserves spatial information and templates rapid resumption of morphogenesis upon reanimation. In *C. elegans* embryos, anoxia drives progressive assembly of the polarity protein PAR-3 into solid-like clusters that preserve the polarity axis during metabolic arrest despite inactivation of patterning reactions normally required to maintain PAR asymmetry. Embryos expressing cluster-defective PAR-3 fail to maintain asymmetry during arrest and consequently exhibit polarity axis defects upon reanimation, demonstrating that arrested PAR-3 clusters function as physical templates for re-establishment of polarity. Our data suggest that cells cope with transient metabolic arrest by reversibly converting actively maintained biochemical patterns into stable physical templates that preserve spatial information for later reactivation.

## Introduction

Living systems are able to maintain themselves at states that lie far from equilibrium by harnessing the flow of matter and energy from their environments (Prigogine, 1978; Schrödinger, 1944). Yet organisms have also evolved an ability to enter states of suspended animation, dormancy, or diapause to cope with variable environments and intermittent reduction in resource availability, be that nutrients, water, light, or oxygen (Bickler and Buck, 2007; Buck and Hochachka, 1993; Maire et al., 2020). In such energy-limited states, physiological functions are slowed or arrested altogether, curtailing energetic demands to maintain viability until the return of favorable conditions (Fenelon et al., 2014).

Adaptation to suspended animation involves rapid downregulation of energy intensive activities such as protein translation, as well as extensive metabolic and physical remodeling (Rosswag De Souza et al., 2025). One common feature of such remodeling includes the formation of molecular assemblies or condensates, including stress granules, proteasome storage granules, polymers of metabolic enzymes, or even wholesale solidification of cytoplasm (Boothby et al., 2017; Franzmann and Alberti, 2019; Heimlicher et al., 2019; Helena-Bueno et al., 2024; Joyner et al., 2016; Lynch et al., 2020; Munder et al., 2016; Parry et al., 2014; Protter and Parker, 2016; Schisa, 2014; Tang et al., 2026). These transitions dramatically reshape the activity and dynamic properties of the constituent molecules, in some cases sequestering key molecules and maintaining them in inactive, protected or “poised” states that can be rapidly redeployed when favourable conditions return. It has even been suggested that condensates can serve as a form of cellular memory (Maity and Moschou, 2026).

Less is known about how cells preserve spatiotemporal information required to resume normal function. This poses a particular challenge to embryos as development relies on active self-organizing processes to generate and maintain the spatial cues that specify form, fate and function (Goldbeter, 2018; Kirschner et al., 2000). Because these systems are maintained in a far-from-equilibrium state by ongoing biochemical activity, collapse of energy-dependent processes would be expected to destabilize spatial organization. How spatial information nevertheless persists during energetic collapse remains unclear.

Cell polarity provides spatial asymmetries that guide morphogenesis in early metazoan development (Campanale et al., 2017). In many embryonic systems, the PAR polarity network plays a central role, integrating intra- and extracellular cues to generate properly oriented polarized membrane domains that template downstream processes such as spindle positioning, inheritance of cell fate determinants, and acquisition of polarized morphologies (Goldstein and Macara, 2007). Both the segregation and maintenance of asymmetric polarity domains rely on active biochemical patterning processes (Goehring, 2014; Lang and Munro, 2017). Thus, polarity reflects precisely the type of active information-bearing system predicted to be vulnerable to metabolic arrest.

*C. elegans* embryos are particularly dependent on PAR polarity to guide early morphogenetic events. Throughout early blastomere divisions, PAR proteins control embryonic axis specification, segregation of fate determinants, and mitotic spindle positioning such that blastomeres acquire the correct size, fate, and position within the embryo, paralleling polarity-dependent patterning processes in the early mammalian embryo (Delattre and Goehring, 2021; Korotkevich et al., 2017; Nance et al., 2003; Rose and Gonczy, 2014; Stolpner et al., 2023).

In response to oxygen deprivation (anoxia), a condition thought to be common in the environments that constitute their typical habitat, *C. elegans* embryos enter a reversible state of suspended animation (Padilla and Ladage, 2012; Van Voorhies and Ward, 2000). This arrested state is characterized by cessation of cell cycle progression, dramatically reduced ATP:ADP ratios, and changes in chromatin architecture (Hajeri et al., 2005; Padilla et al., 2002). Arrested embryos remain viable for up to several days and resume development upon restoration of normal levels of environmental oxygen (Padilla et al., 2002). Thus, *C. elegans* embryos present a highly tractable system to ask whether cells encode a stable, energy-independent spatial template of the pre-arrested state that can be used to re-establish spatial organization upon reanimation.

While PAR polarity is guided by specific spatiotemporal cues (Anderson et al., 2008; Cowan and Hyman, 2004; Munro et al., 2004; Ng et al., 2023; O’Connell et al., 2000; Wallenfang and Seydoux, 2000), the patterns of PAR protein localization that define the polarity axis arise via a dynamic process of self-organization (Goehring et al., 2011b; Gross et al., 2019; Motegi et al., 2011; Sailer et al., 2015). In the well-studied P lineage blastomeres, mutually antagonistic groups of anterior and posterior PAR proteins partition into opposing membrane domains via energy-dependent cycles of phosphorylation and GTP hydrolysis (Figure 1A). Most critically, PKC-3 kinase activity excludes posterior PAR proteins from the anterior membrane, while posterior proteins PAR-1 and CHIN-1 exclude anterior PAR proteins from the posterior (Beatty et al., 2013; Benton and St Johnston, 2003; Betschinger et al., 2003; Boyd et al., 1996; Calvi et al., 2022; Etemad-Moghadam et al., 1995; Gotta et al., 2001; Hoege et al., 2010; Hurov et al., 2004; Kumfer et al., 2010; Moreira et al., 2019; Tabuse et al., 1998; Traweger et al., 2008; Watts et al., 1996).

**Figure 1:**
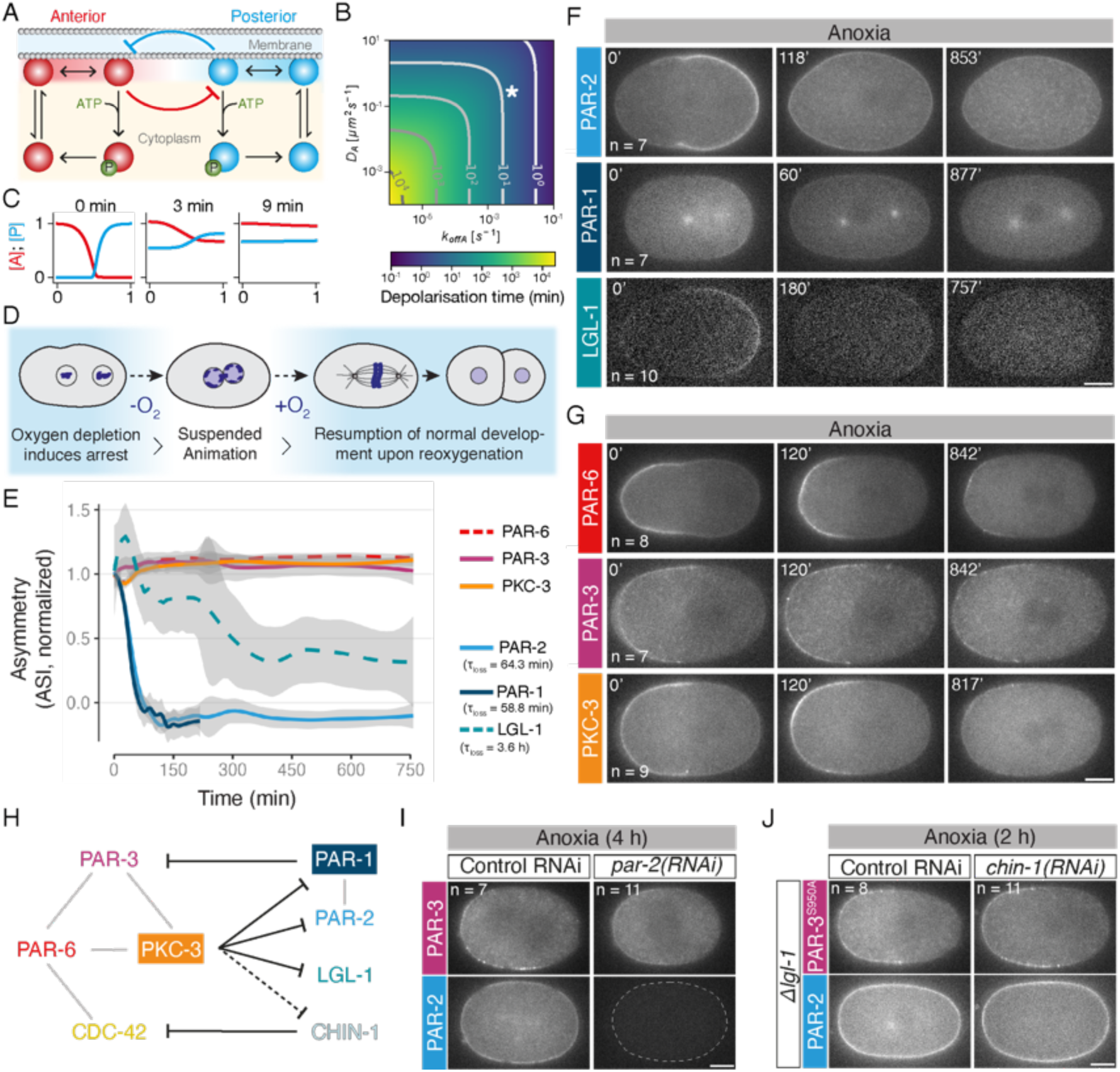
Decoupling of anterior and posterior PAR protein localization during suspended animation. **A.** Schematic representation of a mutual antagonism model for PAR polarity (i.e. (Goehring et al., 2011b)). PAR proteins reversibly bind to the plasma membrane and diffuse laterally at the plasma membrane. PAR polarity domains are maintained by mutually antagonistic phosphorylation reactions: Anterior (“A”, red) and posterior (“P”, blue) PAR proteins phosphorylate each other, which promotes membrane dissociation. Molecules that dissociate from the membrane enter a rapidly mixed cytoplasmic pool and become available to rebind the membrane at any position. Thus, polarized distributions of PAR proteins require continuous active membrane displacement via phosphorylation. As anoxia is expected to limit ATP, one would expect phosphorylation-dependent mutual antagonism to be compromised leading to loss of polarity. **B.** For a simple mutual antagonism model as in (**A**), once the stabilizing cross-phosphorylation reactions cease, polarity decays as a function of the diffusivity on the cell membrane (*D*) and the rate of exchange with the well mixed cytoplasmic pool (*k*_off_)). For an indicated parameter set (*D* = 0.28µm^2^/s, *k*_off_ = 0.0054s^−1^) approximating what has been reported for bulk turnover of PAR-6 (Goehring et al., 2011a; Robin et al., 2014), polarity decays within minutes. **C.** PAR distributions for the indicated parameter set (* in **B**) after 0, 3 and 9 minutes highlight rapid decay of asymmetry. **D.** Schematic of anoxia-induced arrest. Zygotes are mounted in oxygen-scavenging buffer to induce anoxia, resulting in arrest within ∼20 minutes (−0_2_, white region). Zygotes can be maintained in a reversible, arrested state in anoxia for up to 3 days, though viability declines dramatically beyond 24 hrs. Upon re-oxygenation (+0_2_), zygotes re-enter the cell cycle and resume normal development, including undergoing a normal asymmetric division. **E-G.** Selective maintenance of polarity of anterior, but not posterior PAR proteins during anoxia**. (E)** Asymmetry index (ASI) shown for the indicated PAR proteins during the transition to anoxia, normalized to the distribution of the respective proteins at the onset of anoxia (mean ± 95% CI). Fitting of decay curves to a simple exponential yields the timescale of polarity loss (**τ**_loss_). Example midplane confocal images corresponding to quantification in (E), showing GFP-pPAR **(F)** and GFP-aPAR fusions **(G)**, *n* indicated. For consistency, analysis was restricted to embryos that arrested between pronuclear meeting and metaphase (maintenance phase). ASI is a signal-normalized measure of the difference in intensity at the anterior and posterior poles and defined as ASI = |I_A_ - I_P_| /(I_A_ + I _P_), here shown normalized to the value at time = 0 min, such that ASI = 1 when protein is polarized, 0 when uniform. Images showing localizations of aPAR proteins at the cortical plane in anoxia can be found in Figures 2A, S2B. Movie S1 shows timelapse of PAR-2 and PAR-3 during entry to anoxia. **H.** Schematic of antagonistic interactions in the core of the PAR network. PKC-3 and PAR-1 are the anterior and posterior kinases, respectively. aPAR asymmetry is thought to be maintained via phosphorylation of PAR-3 by PAR-1, and GAP activity of CHIN-1 which locally suppresses CDC-42 activity. How LGL-1 maintains aPAR polarity is less clear. Inhibitions by phosphorylation are represented in black; grey lines correspond to known physical interactions; dashed lines are used to indicate unknown links. See recent reviews (Goehring, 2014; Lang and Munro, 2017; Motegi and Seydoux, 2013). **I-J.** Maintenance of aPAR polarity in anoxia is not affected either by depletion of PAR-2 **(I)** or combined disruption of known pPAR inhibitory pathways from LGL-1, PAR-1, and CHIN-1 **(J)**. Number of embryos *(n*) indicated. PAR-3^S950A^ lacks the dominant PAR-1 phosphorylation site and can not be actively excluded from the posterior by PAR-1 (Motegi et al., 2011). Dashed line in (I) indicates embryo boundary highlighting depletion of PAR-2 signal. PAR-3::GFP, PAR-2::mCh expressing lines used. All scale bars, 10µm.

**Figure S1 (Figure 1):**
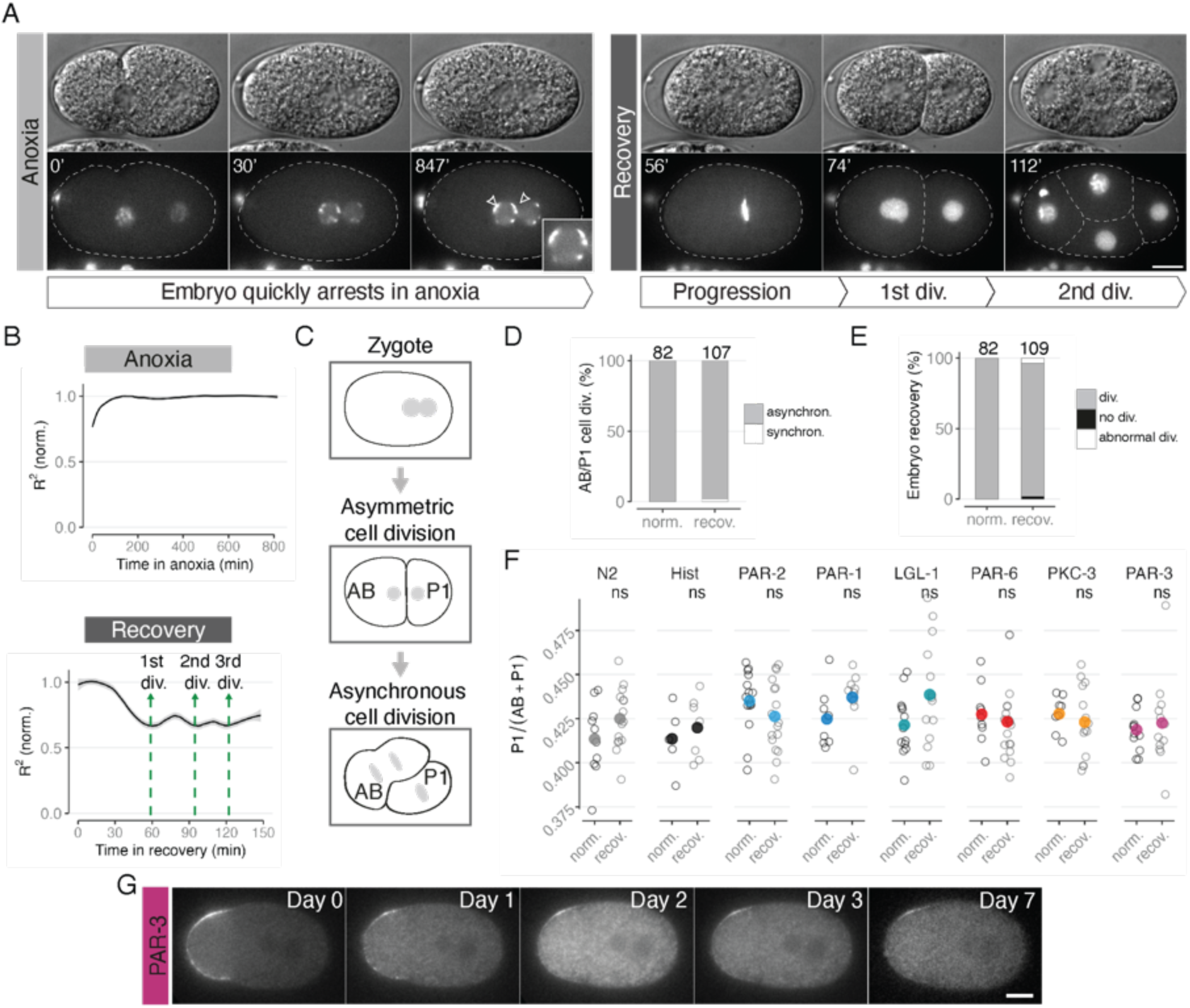
Rapid arrest and recovery of zygotes subject to transient anoxia. **A.** Rapid induction of anoxia and recovery from anoxia in zygotes using wash-in/out of oxygen scavenging buffers. Midsection DIC and confocal images of mCh:Histone highlight anoxia arrest characterized by association of chromosomes with the nuclear periphery (white arrows). Inset shows a magnification of one of the pronucleus. Upon recovery, the cell cycle resumes, chromosomes congress and embryos undergo normal cell division. Dashed lines outline individual blastomeres. Time (min) shown relative to anoxia onset (t0). **B.** Arrest is accompanied by cessation of yolk granule movement. This can be quantified by measuring the correlation coefficient (R^2^) between frames, which progressively reaches 1 upon full arrest (top). Once oxygen is reintroduced to the environment (recovery), the zygote reanimates, which is associated with a decrease in R^2^ (below). Note, the cyclic R^2^ drops during recovery correspond to heightened morphological changes during cell division events. **C.** Schematic of normal asymmetric division highlighting the cell size asymmetry and cell cycle asynchrony between AB and P1 daughters of the zygote. **D.** Fraction of embryos exhibiting asynchronous (asynchron.) or synchronous (synchron.) divisions for a regime in which embryos never experienced anoxia (normoxia, “norm.”) or a regime in which embryos experienced overnight anoxia and were then recovered (recovery, “recov.”). Asynchrony is defined as AB dividing before P1. Note less than 2% of embryos show synchronous divisions after anoxia recovery. Pooled data from the different lines is shown, number of embryos (*n*) indicated. **E.** Fraction of embryos undergoing normal division following recovery from anoxia. An embryo was considered “recovered” if it underwent two division cycles (div.) during the 2 h following recovery (which guarantees enough time for the one-cell embryo to become a 4-cell embryo). If embryos do not divide during this period, they are classified as “no div.”. Abnormal division (“abnormal div.”) corresponds to situations in which either the polar body was clearly re-absorbed during recovery or when P1 and AB cells divided synchronously. Pooled data from the different lines in (F) is shown, number of embryos (n) indicated. 75.77% ± 7.46 (N2) and 85.5% ± 9.54 (PAR-3 GFP) embryos hatched into viable larva. **F.** Daughter size asymmetry (here measured by relative P1 size) is unaffected in embryos after recovery from anoxia. P1 size shown for embryos never experiencing anoxia (norm) and matched embryos dividing after recovery from anoxia for a range of worm lines. Individual embryos (open circles) and mean values shown (closed circles, color-coded by protein). Embryos arresting between late establishment and metaphase were used. **G.** PAR-3(GFP) asymmetry is maintained for extended periods in anoxia (see also Figure 3a (bottom panel) for images taken at the cortical plane of a similarly arrested embryo). All scale bars, 10µm.

Consistent with this model, acute inhibition of PKC-3 leads to the collapse of polarity domains and the loss of asymmetry in downstream pathways that are critical for asymmetric division (Hannaford et al., 2019; Ng et al., 2022; Rodriguez et al., 2017). Thus, embryonic cell polarity represents a non-equilibrium, energy-consuming steady state, whose spatial organization is predicted to collapse when ATP becomes limiting (Cross and Hohenberg, 1993; Goehring et al., 2011b; Khuc Trong et al., 2014) (Figure 1A-C). Intriguingly, polarity is disrupted during hypoxia in both developmental and disease contexts, supporting conserved links between oxygen restriction and polarity maintenance (Dong et al., 2015; Hapke and Haake, 2020; Lu et al., 2021).

Here we show that the asymmetry of many polarity components was lost as *C. elegans* embryos entered anoxia-induced suspended animation. However, the core anterior PAR proteins PAR-6, PKC-3, and PAR-3 remained asymmetric through their incorporation into hyper-stable membrane-associated clusters of PAR-3. These assemblies remained asymmetric throughout arrest and were required to properly restore polarity domains upon reanimation. Thus, anoxia-induced changes in polarity protein dynamics allow cells to physically encode a persistent, energy-independent memory of the embryonic polarity axis during suspended animation, thereby ensuring that embryos can rapidly resume normal development once favorable conditions return.

## Results

### Suspended animation selectively preserves aPAR polarity

To monitor how PAR polarity behaves during anoxia-induced suspended animation, we developed a system to rapidly and reversibly deplete oxygen from embryos (Figure 1D, Figure S1A). Mounting embryos in oxygen scavenging buffer based on protocatechuic acid/protocatechuate-3,4-dioxygenase (PCA/PCD) (Aitken et al., 2008) rapidly induced developmental arrest within 30 min, characterized by cessation of cell cycle progression, peripheral chromosome positioning, and reduced cytoplasmic movements (Foe and Alberts, 1985; Hajeri et al., 2005). Cellular organization did not change over the ensuing 12+ hours (Figure S1A-B). Reoxygenation triggered reanimation with a distinct lag, followed by resumption of cytoplasmic movements, asymmetric divisions, and normal embryonic development, with more than 94% of embryos dividing normally through two subsequent cell divisions and >75% of embryos hatching into larvae (Figure S1A-F).

This setup allowed us to examine the response of both anterior (aPAR) and posterior PAR (pPAR) proteins during anoxia. Consistent with loss of active polarity maintenance, all pPAR proteins lost their normal pattern of localization following oxygen depletion, though with distinct dynamics and final states (Figure 1E-F). Within one hour, both PAR-1 and PAR-2 expanded into the anterior and became symmetrically distributed, with PAR-1 eventually becoming undetectable at the plasma membrane. By contrast, posterior membrane localization of LGL-1 gradually decreased, becoming undetectable by 3 hours. While the basis for these differences is unclear, these results demonstrate that the active mechanisms for maintaining pPAR polarity collapse during suspended animation.

Despite loss of pPAR asymmetry, we found that anterior PAR proteins (aPARs), including PAR-3, PAR-6, and PKC-3, remained asymmetric throughout suspended animation (Figure 1E, 1G). This asymmetry persisted well beyond the time (≥ 3 days) at which embryos have been reported to lose viability (Figure S1G). The asymmetry of aPARs in anoxia was not dependent on pPAR proteins as it was retained normally in embryos depleted for PAR-2, or lacking PAR-1, LGL-1, and CHIN-1 in combination (Figure 1H-J). Thus, in contrast to expectations of current theoretical understanding of PAR polarity maintenance, aPAR and pPAR localizations become uncoupled during anoxia such that aPARs retain asymmetry despite loss of pPAR polarity.

### Persistent aPAR asymmetry is associated with PAR-3-dependent clusters

Given that the principles governing polarity maintenance change during anoxia, we examined the behavior of aPAR proteins in more detail. Under normal conditions, PAR-3 localizes to dynamic membrane-associated clusters, while PAR-6/PKC-3 heterodimers cycle between a PAR-3-associated clustered state and a rapidly diffusing CDC-42-associated state (Dickinson et al., 2017; Rodriguez et al., 2017).

We found that PAR-3 clusters were not only maintained during the transition into anoxia, but became more prominent as cells arrested, with cluster number and size increasing over the first hour in anoxia before plateauing (Figure 2A-C, S2A). While some clusters appeared in the posterior, increased PAR-3 recruitment was largely restricted to the anterior, reinforcing the initial asymmetry (Figure 2D-E). Coincident with increased PAR-3 clustering, PAR-6 and PKC-3 became visibly enriched in clusters at the expense of the diffuse state (Figure 2A-C). This enrichment was accompanied by increased colocalization of PAR-6 with PAR-3 and was dependent on PAR-3 (Figure 2F-G, Movie S3). By contrast, clustering was, if anything, enhanced further upon depletion of CDC-42, consistent with an anoxia-induced shift of PAR-6/PKC-3 towards the PAR-3-associated state (Figure 2H-I).

**Figure 2:**
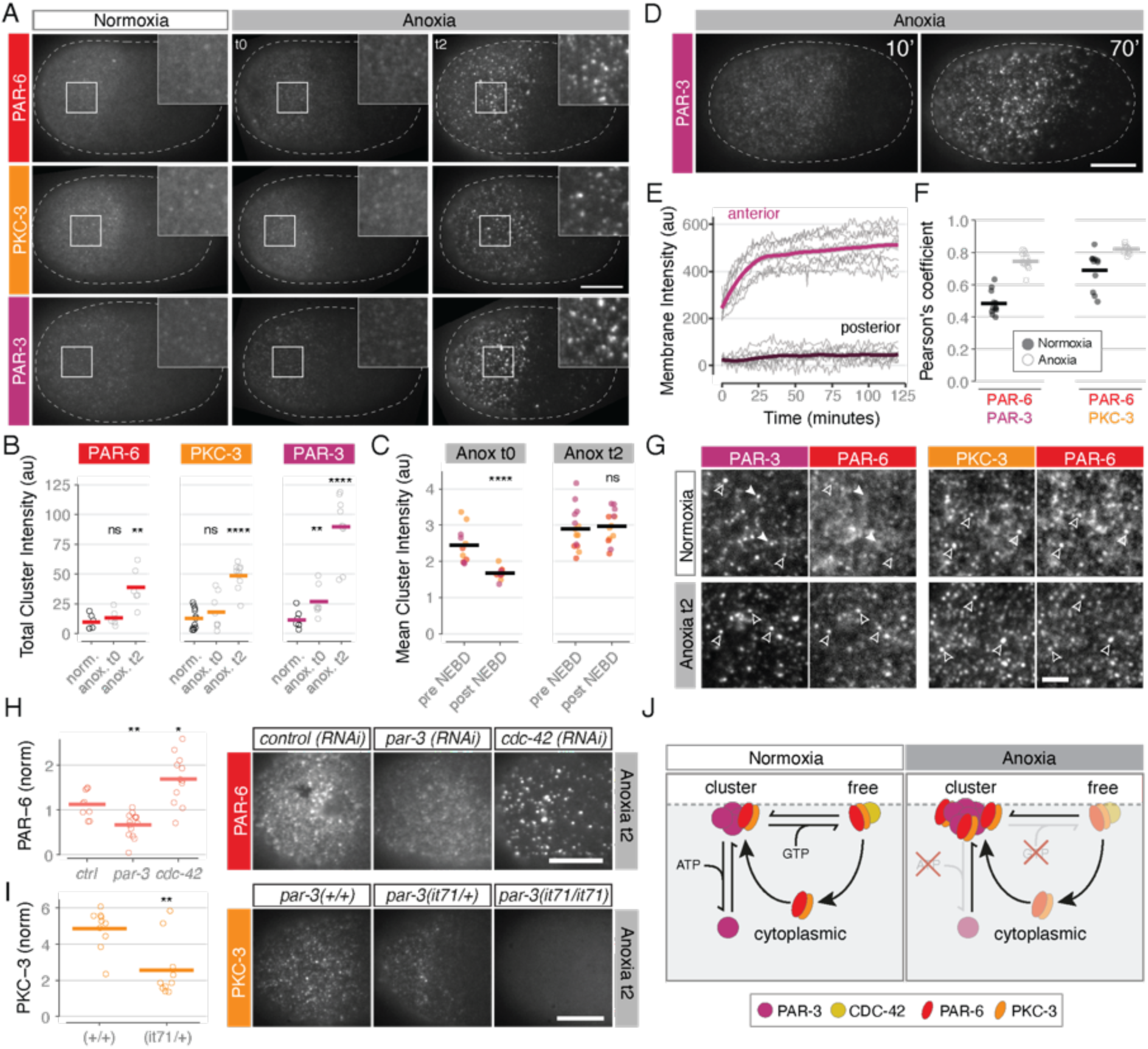
aPAR asymmetry is associated with persistent PAR-3-dependent clusters. **A.** Anoxia enhances aPAR clustering. Example cortical images of the indicated PAR proteins shown in normoxia, at anoxia entry (anox. t0) and after 2h in anoxia (anox. t2) for post-NEBD embryos. Example of pre-NEBD embryos are shown in Figure S2B. Insets show 2-fold magnification. Dashed lines indicate embryo boundary. All lines are GFP fusions. **B.** Quantification of area-normalized integrated cluster intensity for embryos in normoxia, at the entry to anoxia (t0) and after 2 hrs in anoxia (t2). Data for individual embryos (open circle) and means (lines) shown. (ns: p > 0.05, *: p <= 0.05, **: p <= 0.01, ****: p <= 0.0001, Wilcoxin test w/ normoxia as reference). **C.** Embryos arrested pre- or post-NEBD achieved similar PAR-3 cluster sizes in anoxia, although aPAR cluster sizes were initially smaller in post-NEBD embryos at t0 due to cell cycle regulation of PAR-3 oligomerization. Quantitation of mean cluster size (arbitrary units) showing mean (line) and individual embryos. Note data for all aPARs pooled, with individual embryos colour-coded by protein (see Figure 2B for colour reference). (ns: p > 0.05, ****: p <= 0.0001, Wilcoxin test w/ pre-NEBD as reference). **D-E.** PAR-3 undergoes preferential accumulation in the anterior during anoxia. (D) Cortex images of cluster behavior showing accumulation of clusters in anterior. Dashed lines indicate embryo boundary. (E) Quantification of protein accumulation over time from midplane images. Individual embryo data shown along with means. PAR-3::GFP shown. **F-G**. Anoxia drives redistribution of PAR-6 into PAR-3 clusters. Sample images (G) and quantification of colocalization of PAR-6::mCh and PAR-3::GFP (F) shown in normoxia and in anoxia. Colocalization of PAR-6::mCh and PKC-3::GFP, which are thought to form a stable heterodimer, are shown for reference. Empty arrowheads show areas of colocalization between aPARs, filled arrowheads show areas in which clusters do not co-localise.Mean (closed circle) and individual embryo (open circles) indicated. See Movie S3 for timelapse. **H-I.** PAR-6- and PKC-3-positive clusters are dependent on PAR-3. Sample images (left) and quantification of total, area-normalized, intensity for PAR-6::GFP(H) and PKC-3::GFP(I) for maintenance phase embryos under the indicated conditions at anoxia t2. Note total cluster intensity is reduced and enhanced when PAR-3 and CDC-42 levels are reduced, respectively. Individual (open circle) and mean (closed) values shown. (*p <= 0.05, **p <= 0.01, Wilcoxon test vs control as reference). “par-3” and “cdc-42” correspond to par-3- and cdc-42-RNAi, respectively. The *it71* allele harbours a maternal effect lethal mutation that prevents PAR-3 expression in the germline and early embryo. +/+, it71/+, and it/71/it71 indicate wild type, heterozygous and homozygous *it71*. **J.** PAR-6 and PKC-3 shuttle between distinct states, as shown previously (Rodriguez et al., 2017): (1) a free cytoplasmic state, (2) a clustered, plasma membrane- and PAR-3-associated state, and (3) a diffuse plasma membrane- and CDC-42-associated state. In normoxia, energy-dependent processes drive flux out of the clustered state resulting in a balance of aPARs between clustered and non-clustered states (left). In anoxia, a reduction in these energy-dependent processes leads to a shift in aPARs towards the PAR-3 clustered state (right). All scale bars, 10µm.

**Figure S2 (Figure 2):**
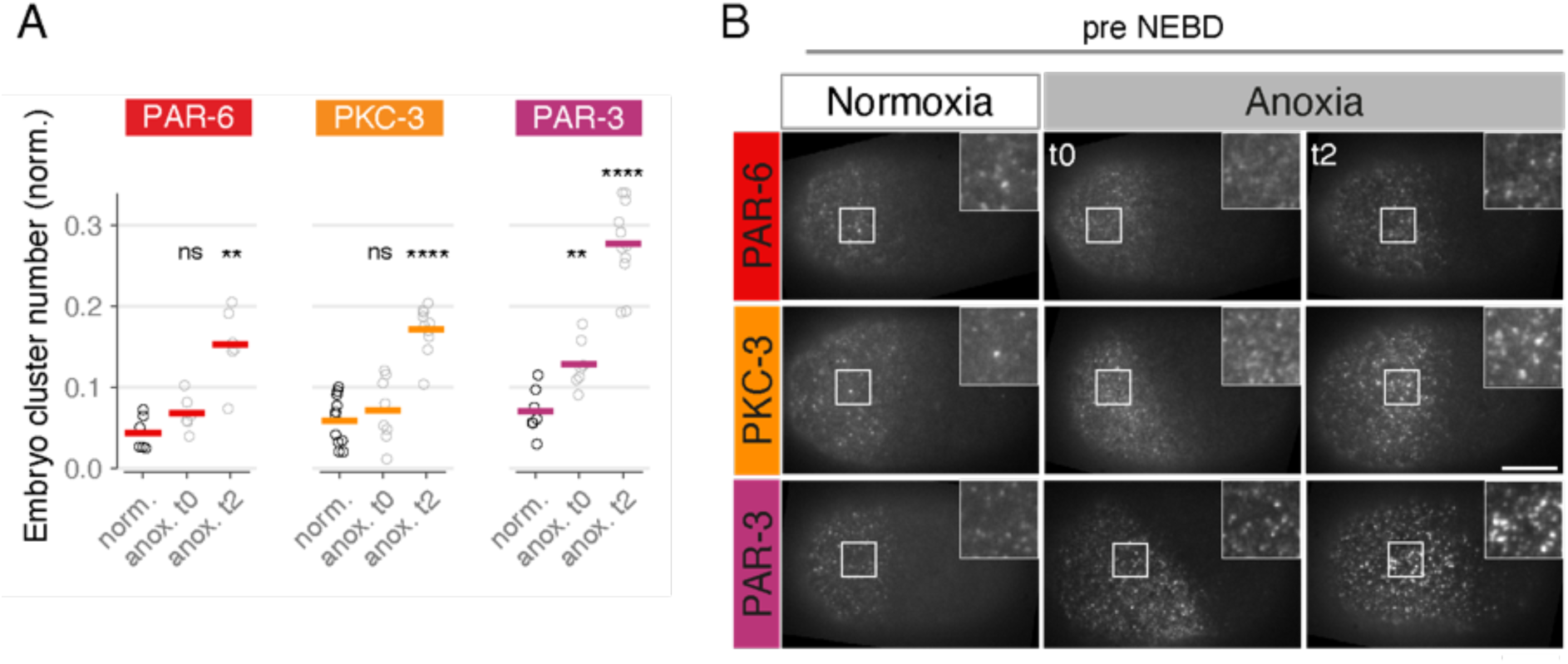
aPAR asymmetry is associated with persistent PAR-3-dependent clusters, continued. **A.** Quantitation of area-normalized integrated cluster number for embryos in normoxia (norm.), at the entry to anoxia (anox.) (t0) and after 2 hrs in anoxia (t2). Data for individual embryos (open circle) and means (lines) shown. (ns: p > 0.05, *: p <= 0.05, **: p <= 0.01, ****: p <= 0.0001, Wilcoxin test w/ normoxia as reference). GFP fusions shown. **B.** Additional examples of anoxia-induced clustering of aPARs for pre-NEBD embryos. Note aPARs are initially more clustered than post-NEBD embryos at t0 (see Figure 1A, 1C), but otherwise the pattern is similar. Representative embryos for each condition are shown. Insets show a magnification of the indicated area. Anoxia t0 corresponds to anoxia start, t2 corresponds to 2 h in anoxia. Scale bar is 10 µm.

Because PAR-3 clustering is regulated by the cell cycle via PLK-1 (Dickinson et al., 2017), we wondered whether the maintenance of asymmetry depended on the stage of cell cycle arrest. In normal conditions, clustering increases steadily throughout the polarity establishment phase before being suppressed in the polarity maintenance phase, a period spanning the time of pronuclear meeting to anaphase (See also (Dickinson et al., 2017; Wang et al., 2017)). Despite initial differences in clustered PAR-3 at entry to anoxia (t0), embryos converged to similar distributions of PAR-3 by 2 hours (t2) regardless of the cell cycle stage at arrest (Figure 2C, 2A vs S2B). This observation suggests that embryos enter a common arrested polarity state regardless of when oxygen is removed.

We conclude that, rather than aPARs simply being maintained in their normal distribution, anoxia remodels the anterior polarity network with PAR-6 and PKC-3 consolidated into asymmetric PAR-3-dependent clusters. While the mechanism driving this change is unclear, it is tempting to speculate that this reorganisation emerges naturally from the existence of two active mechanisms: one that directly suppresses PAR-3 oligomerization via PLK-1 activity (Dickinson et al., 2017) and one that promotes conversion of PAR-3-associated PAR-6/PKC-3 complexes into the active, more diffusive CDC-42-associated state via PKC-3 activity (Rodriguez et al., 2017). Such a system would naturally relax into a more stable clustered state upon energy withdrawal as these active processes slow (Figure 2J, Normoxia vs. Anoxia).

### PAR-3 clusters undergo progressive immobilization in anoxia

Given this enhancement of asymmetric PAR-3 clusters, we next asked what features of PAR-3 clusters could drive long-term persistence of asymmetry. In normoxia, clusters are highly dynamic. They exhibit lateral motion on the membrane, and form and dissipate over minutes to tens of minutes (Dickinson et al., 2017; Rodriguez et al., 2017; Sailer et al., 2015; Wang et al., 2017). In addition, aPARs rapidly exchange into and out of clusters on the seconds-to-minutes timescale (Figure 3C) (Chang and Dickinson, 2022; Illukkumbura et al., 2023; Lang et al., 2026). Together, such dynamics would be expected to destabilize asymmetries, particularly over the extended timescales of anoxia-induced arrest. This led us to examine how these dynamics may change in anoxia.

**Figure 3:**
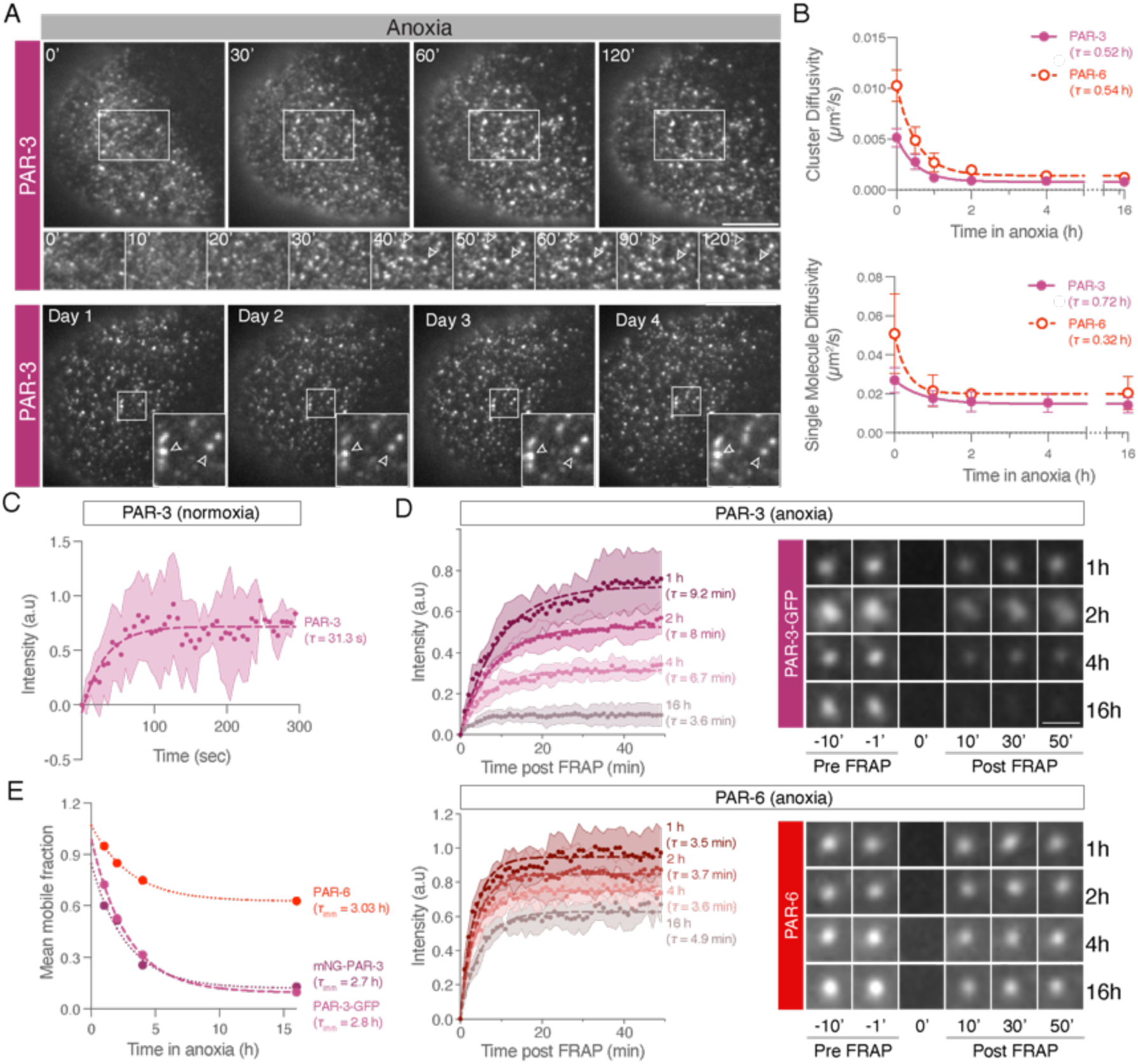
PAR-3 undergoes progressive immobilization during suspended animation. **A.** PAR-3(GFP) clusters undergo progressive immobilization in anoxia (top) and remain immobile for days (bottom). Cortical plane (TIRF) showing only the anterior domain of embryos. Top panels: time in minutes, with additional insets below. Bottom panels: time in days, insets magnified. Arrowheads show immobile clusters. Scale bar is 10 µm. **B.** PAR-3(GFP) and PAR-6(GFP) diffusion decreases during anoxia entry, shown at the level of clusters (left) and single molecule (SM) (right). Curves were fit to a one-phase decay curve; respective timescales and error bars shown. **C.** PAR-3 clusters are dynamic in normoxia. Fluorescence recovery curves after photobleaching plotted with mean (circles), error (shaded region), and fit (dashed line) shown. Bleaching occurred at “0 min”. Data normalized to initial pre-bleach fluorescence and fit to single component exponential to yield characteristic turnover time, *τ*. Time (sec). **D.** Turnover of PAR-3 (top) and PAR-6 (bottom) after FRAP after 1h, 2h, 4h, 16h in anoxia. Pre-bleach normalized fluorescence recovery shown with time series images of representative clusters subject to FRAP. Time(min) with first post-bleach frame at time = 0 min. Mean (circles), error (shaded region), and fits to a one-component exponential recovery (dashed line) shown, along with the best fit value for the recovery timescale, *τ*. Note that *τ* is relatively unchanged in anoxia, suggesting that long-term anoxia primarily drives a switch between mobile and immobile states. Scale bar is 1µm. **E.** Progressive immobilization of PAR-3 during anoxia. Plot of mean mobile fraction vs anoxia duration for PAR-3 and PAR-6. Mean values obtained from fits of fluorescence recovery for individual embryos after 1, 2, 4, or 16 h in anoxia shown (closed circles) along with fits to an exponential decay model (dashed lines) to obtain the characteristic immobilization timescale, *τ*_imm_. Sample images shown at right. Note that PAR-3 ultimately exhibits complete immobilization during anoxia, while PAR-6 retains a >0.6 mobile fraction after 16 h in anoxia. Note that similar results were obtained for PAR-3::GFP and mNG::PAR-3.

In anoxia, clusters became larger and more stable, reaching a stable configuration within the first 1-2 hrs of oxygen depletion (Figure 3A, top). Thereafter, clusters were stable not only over the course of hours, but days (Figure 3A, bottom). Tracking of individual clusters revealed that the effective diffusivity of clusters abruptly declined within the first hour of anoxia, reaching ∼10^−4^µm^2^/s, which is likely an overestimate given limits in positional accuracy under our imaging conditions (Figure 3B).

Single-molecule tracking of GFP-fusions to PAR-3 also revealed a reduction in diffusivity. When we examined PAR-6, we found that it was initially more mobile than PAR-3, reflecting its partitioning between clustered and non-clustered states. However, its diffusivity became indistinguishable from that of PAR-3 in anoxia, consistent with its redistribution to a preferentially cluster-associated state (Figure 3B). We conclude that aPAR mobility is dramatically reduced in anoxia, consistent with consolidation of aPARs into large, immobile clusters.

In addition to cluster mobility, we also examined the turnover of molecules within clusters using fluorescence recovery after photobleaching (FRAP). To this end, we bleached individual clusters of PAR-3 either during normoxia or at multiple times after anoxia entry and fit recovery curves to a simple exponential model with an immobile fraction (Figure 3C-E). In normoxia, we found that PAR-3 exchanged into and out of clusters with a characteristic exchange timescale of ∼31 seconds (Figure 3C). Anoxia induced a progressive decrease in the mobile fraction (Figure 3D-E). The response of PAR-6 was similar; however, whereas PAR-3 became almost completely immobile by 16 hrs, the mobile fraction of PAR-6 remained above 0.5 (Figure 3D-E). Fitting the immobile fraction as a function of duration of anoxia, we obtained animmobilization timescale (***τ***_imm_) of ∼ 3 hrs for both proteins (Figure 3E). As the primary change was in the immobile fraction, loss of dynamic exchange likely arises from a transition of molecules into an immobilized, potentially solid-like state rather than uniform slowing of the entire population.

Thus, anoxia is associated with near-complete immobilization of PAR-3 within membrane-associated clusters. This effectively locks in the asymmetric distribution of other aPARs, rendering it independent of the active energy-dependent processes that are normally required to maintain polarity.

### Persistent aPAR asymmetry requires PAR-3 oligomerization and aPAR feedback

Having shown that persistent aPAR asymmetry is associated with consolidation of aPARs into asymmetric, highly stable PAR-3-associated clusters, we next explored the molecular requirements for this consolidation. Membrane association and segregation of PAR-3 in the zygote depend on two processes: (1) oligomerization of PAR-3, which promotes assembly into stable, slowly diffusing clusters at the membrane (Chang and Dickinson, 2022; Dickinson et al., 2017; Illukkumbura et al., 2023; Lang et al., 2026; Li et al., 2010; Rodriguez et al., 2017; Sailer et al., 2015); and (2) potentially cooperative interactions between PAR-3 and other aPAR proteins, which mutually reinforce their membrane recruitment (Hsu and Dickinson, 2026; Hung and Kemphues, 1999; Lang et al., 2026; Tabuse et al., 1998). We therefore asked whether either process was required to stabilize PAR-3 asymmetry in anoxia.

To test whether persistent aPAR asymmetry depended on PAR-3 oligomerization, we analyzed an oligomerization defective PAR-3 variant PAR-3(Δ69-82). PAR-3(Δ69-82) harbors a small deletion in conserved region 1 (CR1), does not form clusters, and fails to accumulate at the plasma membrane in zygotes due to reduced avidity (Li et al., 2010). Consistent with this membrane-binding defect, PAR-3(Δ69-82) fails to accumulate in the anterior in anoxia (Figure 4A, B, Δ69-82). We then examined the response of two modified variants of PAR-3(Δ69-82) in which membrane association is restored either via fusion to a short membrane-targeting peptide (RitC) or via ectopic dimerization by fusion to a dimeric leucine zipper (2mer). Both variants localize to the plasma membrane, segregate normally, and support normal asymmetric cell division, though membrane levels are reduced relative to WT (Illukkumbura et al., 2023). In our assays both were initially segregated into the anterior. However, while dimerization-competent PAR-3(2mer) was able to maintain and even amplify this asymmetry in anoxia, asymmetry of the monomeric PAR-3(RitC) decayed within the first 1-2 hours in anoxia (Figure 4A, B). Thus, oligomerization of PAR-3 appears to be critical for stabilizing and reinforcing asymmetry during the transition into anoxia.

**Figure 4.**
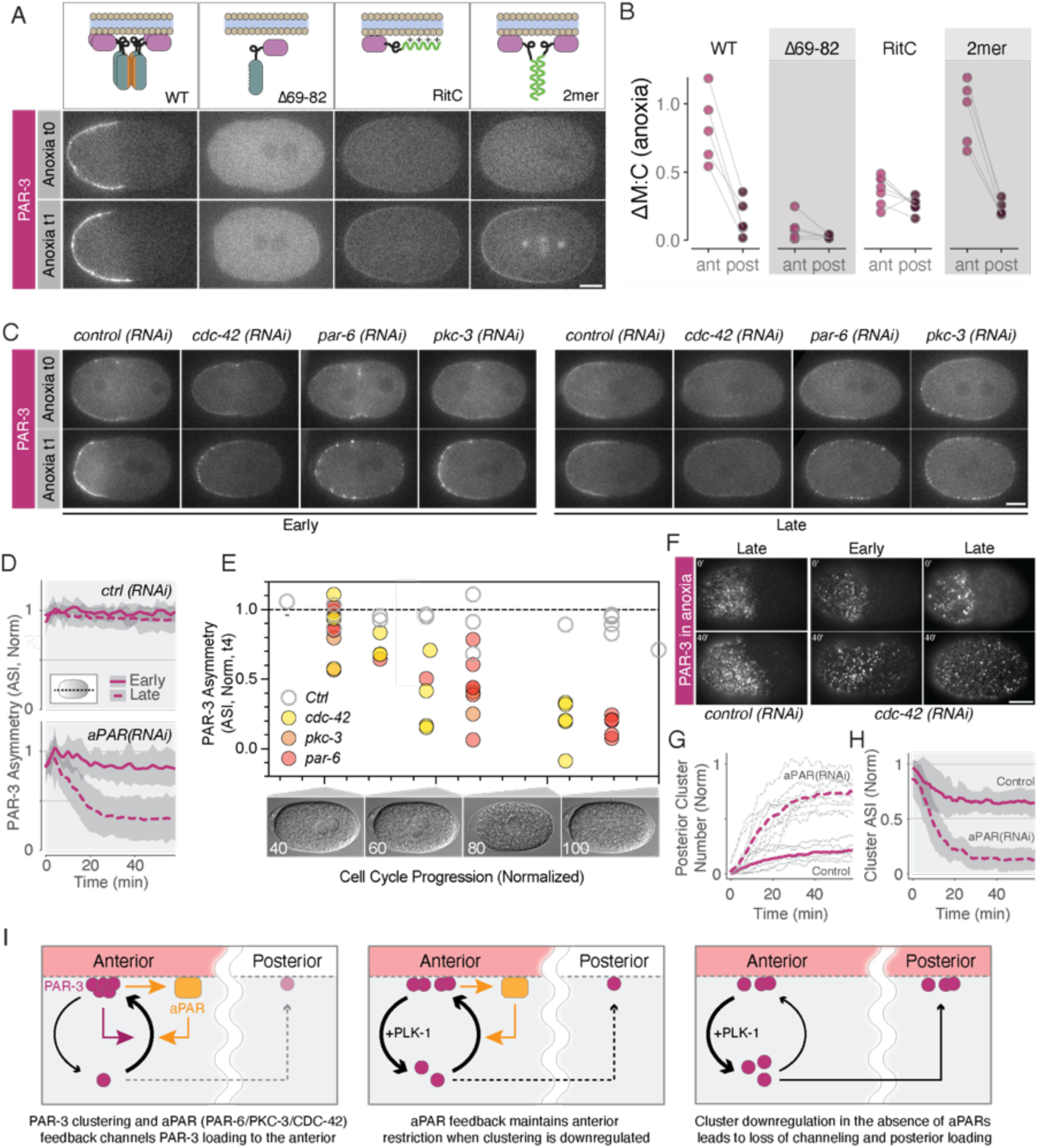
PAR-3 oligomerization and other aPAR proteins contribute to persistent PAR-3 asymmetry in anoxia. **A.** Biased recruitment of PAR-3 during anoxia entry requires oligomerization. Example midplane confocal images of the indicated mNG::PAR-3 variants (upper schematic) shown at the entry to (t0) and after one hour in anoxia (t1). Note only clustered variants (WT, 2mer) maintain stable asymmetry at t1. **B.** Quantification of change in membrane:cytoplasmic (M:C) ratios between t0 and t1 in the anterior (light purple) and posterior (dark purple) for each variant after AF correction for the embryos in (A). Paired data for individual embryos shown (lines). Note only WT and 2mer show preferential accumulation in the anterior. Scale bar, 10µm. **C.** Restriction of PAR-3 to the anterior domain during anoxia entry is stabilized by CDC-42, PAR-6, and PKC-3. Midplane confocal images of PAR-3::GFP expressing embryos subject to the indicated RNAi that arrested either early or late in the cell cycle shown at anoxia entry (0’) and after 40’ in anoxia (∼t1). Scale bar is 10 µm. See (F) for example of images showing cortical plane. **D.** Timecourse of PAR-3 membrane signal asymmetry (ASI) during the transition into anoxia taken from embryos as in (C). Note *cdc-42, par-6, pkc-3(RNAi)* data are pooled - data for individual RNAi conditions in Figure S4. ASI is normalised such that ASI(t0) = 1 for each embryo. Mean ±SD shown. **E.** ASI (PAR-3) after 4h in anoxia (t4) as a function of arrest point in the cell cycle. Cell cycle normalized from 0 (symmetry-breaking) to 100 (anaphase) as judged from DIC images, with 40 ∼ pronuclear meeting, 60 ∼ pronuclear centration, 80 ∼ NEBD, 100 ∼ metaphase. **F.** Depletion of aPARs results in posterior appearance of PAR-3 clusters. Cortical images of PAR-3::GFP embryos arrested in either early or late maintenance (pre NEBD and post NEBD) subject to indicated RNAi shown at 0 and 40 min in anoxia. Note appearance of posterior clusters in *cdc-42(RNAi).* See Movie S4 for timelapse. **G.** Timecourse of area-normalized posterior PAR-3 cluster number for late maintenance arresting embryos subject to aPAR or control RNAi measured from near-TIRF images. Note *cdc-42, par-6, pkc-3(RNAi)* data are pooled. Quantification of individual lines is shown in Figure S4. **H.** Timecourse of cluster number asymmetry (ASI) loss for embryos in (G), normalized to mean ASI in control embryos. **I.** Schematic: PAR-3 clustering and aPAR feedback promote PAR-3 consolidation in clusters in anoxia, restricting the availability of free PAR-3 to bind the posterior membrane and nucleate new clusters. aPAR feedback is sufficient and necessary to limit this posterior nucleation during periods of cluster dissolution by PLK-1 (late maintenance). In the absence of aPARs, cluster dissolution leads to increased free PAR-3 resulting in nucleation of clusters in the posterior and loss of asymmetry during the transition into anoxia.

**Figure S4 (Figure 4):**
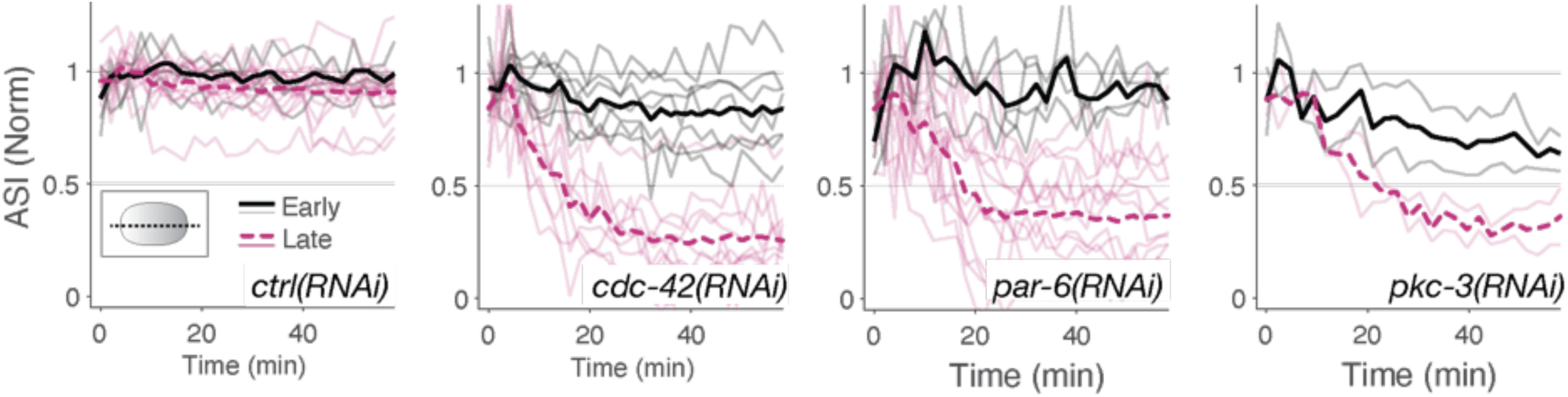
PAR-3 asymmetric accumulation during anoxia entry depends on feedback from other aPAR proteins. Timecourse of PAR-3 membrane signal asymmetry (ASI) during the transition into anoxia (midplane images) of control embryos or embryos depleted of other aPARs that arrest either early or late in the cell cycle. As in Figure 4D, but showing individual embryos traces in RNAi conditions with mean shown.

We next assessed the potential contributions of other aPAR proteins in maintaining asymmetry in anoxia by depleting PAR–6, PKC-3, or CDC-42 via RNAi. In normoxia, depletion of PAR–6, PKC-3, or CDC-42 results in initial segregation of PAR-3, but asymmetry is ultimately lost as clusters are disassembled by increasing PLK-1 activity at mitotic entry (Aceto et al., 2006; Lang et al., 2026; Tabuse et al., 1998). Because PAR-3 remains segregated in PKC-3 inhibited embryos, these proteins likely play a physical rather than enzymatic role in stabilizing PAR-3 asymmetry during cluster disassembly (Rodriguez et al., 2017). We found that maintenance of PAR-3 asymmetry in anoxia depended on all three proteins, but the degree of dependence varied with the timing of arrest (Figure 4C-E). Embryos arresting early in the cell cycle (around the time of pronuclear meeting) retained robust asymmetry upon depletion of PAR-6, PKC-3, or CDC-42. By contrast, embryos that entered anoxia later when PLK-1-dependent cluster disassembly begins (pronuclear centering to NEBD) exhibited substantial to near complete loss of asymmetry. Loss of asymmetry coincided with substantial nucleation of new PAR-3 clusters in the posterior (Figure 4F-H). We therefore conclude that oligomerization of PAR-3 is sufficient to maintain stable aPAR asymmetry in anoxia provided clustering remains favoured. However, once embryos enter the cluster disassembly phase, aPAR feedback becomes essential to suppress posterior accumulation of PAR-3 and asymmetry loss.

### Persistent aPAR clusters promote accurate specification of the polarity axis upon reanimation

Our data so far suggest that immobilization of PAR-3 in clusters locks in aPAR asymmetry during anoxia. We next asked whether embryos use the spatial information encoded by PAR-3 clusters to re-establish normal polarity following reanimation. To test this, we arrested embryos in anoxia for 4 hrs and followed PAR protein behavior during reoxygenation.

The first sign of re-animation in PAR behavior was increased accumulation of PAR-6 and PKC-3 at the plasma membrane (Figure 5A and S5A). Subsequently, pPARs were excluded from aPAR domains to re-establish a complementary posterior domain (Figure 5A, 5B and Figure S5A, S5B). The resulting PAR domains invariably reflected the initial polarity of the embryo that was maintained by aPAR asymmetry during anoxia (Figure 5I - WT). Other methods for inducing anoxia showed similar results (Figure S5C, S5D). These results suggest that persistent aPAR clusters function as a “molecular memory” of zygote polarity during anoxia, which can be used to properly reestablish polarity upon reanimation.

**Figure 5:**
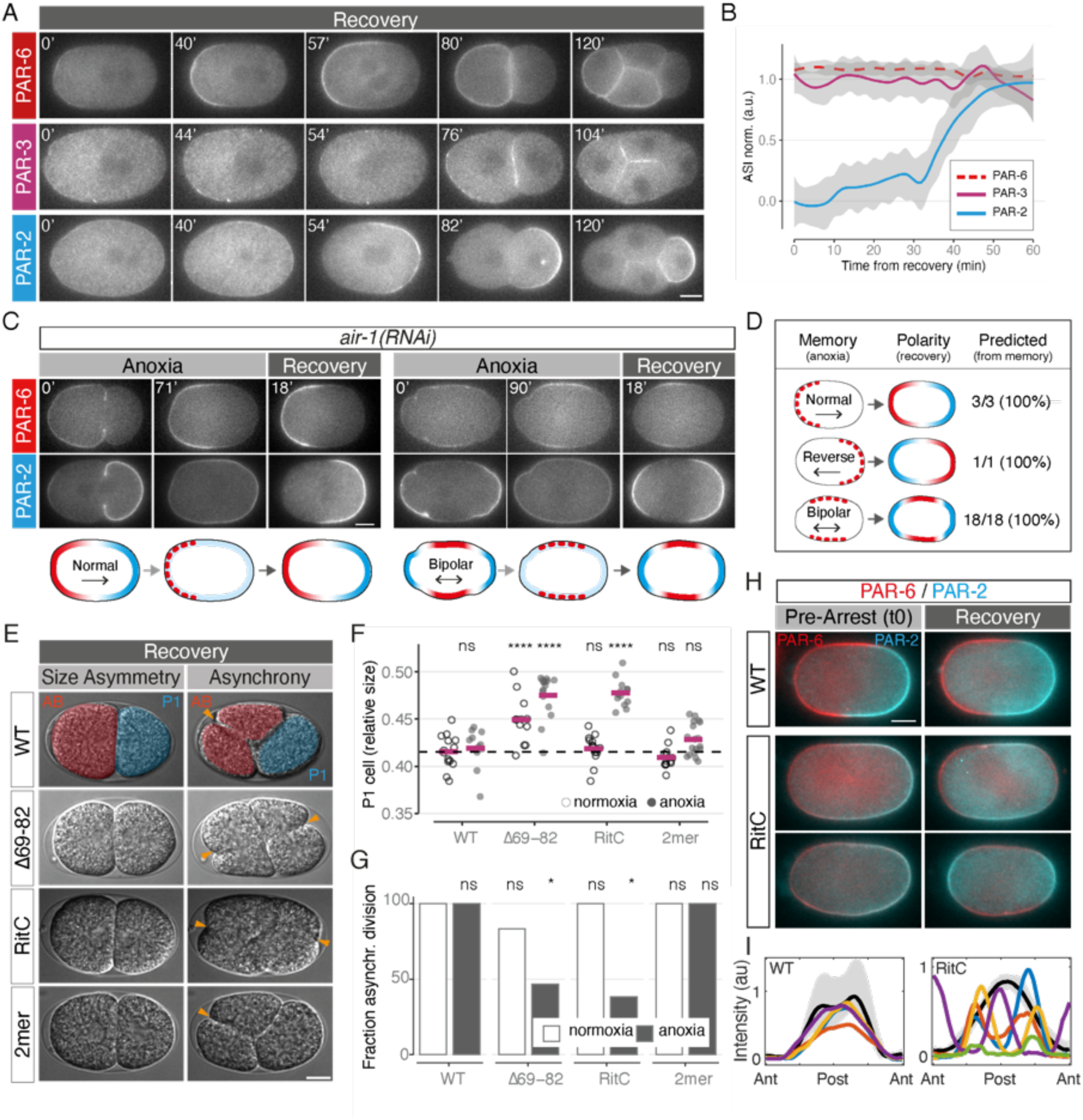
Persistent asymmetric PAR-3 clusters act as a spatial memory which predicts re-establishment of polarity upon recovery. **A-B.** Embryos rapidly restore polarity, including pPAR asymmetry, upon reanimation. Example midplane fluorescence images (A) and quantification of asymmetry (B, ASI) of wild-type embryos expressing PAR-3, PAR-6, or PAR-2 shown during recovery following a period of 4 hrs in anoxia. Time = 0s is the time of buffer exchange to restore normal oxygen levels. Note rapid restoration of PAR-2 asymmetry within 60 minutes of return to normoxic conditions. Smoothed conditional means are shown color-coded for each protein, with respective intervals of confidence. ASI is 1 when protein is polarized, 0 when depolarised. Plasma membrane intensity levels are normalized to the first time point. Data for additional PAR proteins can be found in Figure S5A-B. Similar results were obtained using other methods of anoxia induction (Figure S5C-D). **C-D.** Embryos accurately re-establish PAR distributions from the pre-arrested state. Depletion of AIR-1 results in variable configurations of PAR domains including normal, reversed, and bipolar arrangement of domains. *air-1(RNAi)* embryos expressing PAR-2 and PAR-6 were subject to 4 hrs of anoxia and scored for their ability to restore the pre-arrested PAR distribution based on the PAR-6 “memory.” Example images of normal (left) and bipolar (right) embryos pre-anoxia (0), anoxia arrest (71’/90’), and after recovery (18’) are shown (C), along with the frequency of correct re-establishment of the pre-anoxic pattern of polarity (D). **E-G.** PAR-3 oligomerization is required for normal asymmetric division after anoxia arrest. Embryos carrying the indicated *par-3* alleles were subject to anoxia for 4 hrs and scored for division asymmetry after recovery from anoxia. Asymmetry was scored by size asymmetry (P1/AB+P1) and the presence of division asynchrony between AB and P1. Sample images (E) and quantification (F, G) shown. Note *par-3(RitC)* was specifically defective in division asymmetry after anoxia. In (F), individual (open circles) and mean values are shown. All conditions scored relative to WT, Normoxia (t-test, ns: p > 0.05, ****: p <= 0.0001). **H**. Re-establishment of normal PAR polarity is defective in *par-3(RitC)* embryos, consistent with failure to establish a memory of polarity. WT and *par-3(RItC)* dual PAR-6, PAR-2 labeled embryos were subject to anoxia for 4 hrs and allowed to recover. Midplace confocal images shown at pre-arrest (t0) and following recovery (6 min pre-division). Note variable PAR-2 domain positions in *par-3(RitC)* embryos. A full time course including pre-arrest (Initial, t0), anoxia (t2), and three points during recovery (pre-division, after first division, second division) is shown in Figure S5 along with a normoxia *par-2(RitC)* control for comparison. See Movie S5 for timelapse. **I.** Sample PAR-2 distributions at anoxia entry (grey, mean ±max/min shown) and following recovery (colours show individual embryos) for a set of WT (n=4) and *par-3(RItC)* (n=5) embryos. Note variability of distributions in *par-3(RitC).* All scale bars, 10µm.

**Figure S5 (Figure 5):**
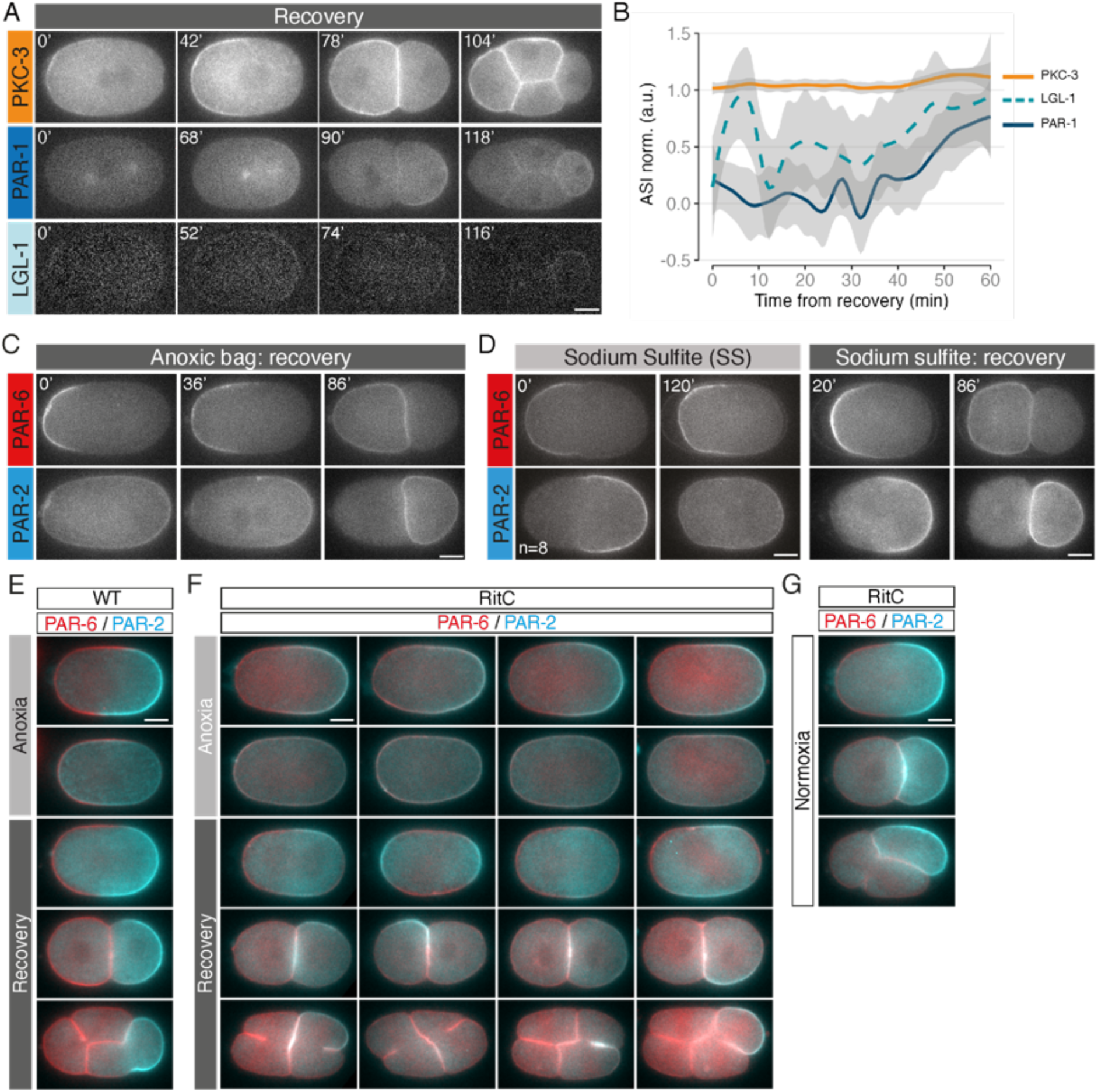
aPAR asymmetry in anoxia is a spatial memory which predicts re-establishment of polarity upon recovery. A-B. Sample images (A) and quantification (B) of PKC-3 (anterior), PAR-1 and LGL-1 (posterior) during recovery following maintenance phase arrest in anoxia. Note these embryos are the same as shown during anoxia in Figure 1E. Start of recovery corresponds to time 0 min. Plasma membrane intensity levels are normalized for the first time point. Note that ASI is 1 when protein is polarised, 0 when uniform. Plasma membrane intensity levels are normalized to the first time point. Smoothed conditional means with confidence intervals shown, colour-coded for each protein. **C.** Sample images of embryos during recovery from anoxia induced by placing worms on plates in an anoxic bag. **D.** Sample images of embryos during entry into and recovery from anoxia induced by mounting in sodium sulfite. **E.** Extended images of control wild-type, dual labeled embryo from Figure 5H showing normal repolarization of PAR-2 and PAR-6 during recovery from anoxia. **F.** Extended images and additional examples of PAR-2 re-polarization phenotypes in par-3(RitC) embryos subject to anoxia, as in Figure 5H (PAR-6, red; PAR-2, cyan). Note embryos 2 and 4 are also shown in Figure 5H. **G.** Sample images of dual labeled *par-3(RitC)* embryos during normoxia at equivalent developmental stages as in anoxia recovery experiments. Note normal size asymmetry and division asynchrony. All scale bars are 10 µm.

To challenge this memory, we depleted embryos of Aurora A kinase, AIR-1. AIR-1 depletion sensitizes the PAR network to alternative polarity cues. *air-1(RNAi)* embryos typically exhibit variable polarization, in which PAR-2 domains can form at either or both poles of the embryo (Kapoor and Kotak, 2019; Klinkert et al., 2019; Reich et al., 2019; Schumacher et al., 1998; Zhao et al., 2019). Using *air-1(RNAi)*, we obtained a mix of normal, reversed, and bipolar embryos (Figure 5C, 5D). In the latter case, PAR-2 was present at both poles with aPARs enriched laterally in a complementary pattern around the middle of the embryo. As in wild-type embryos, aPAR localization in *air-1(RNAi)* embryos was retained in anoxia, while PAR-2 became uniform. Upon recovery, aPAR localisation in anoxia invariably predicted the pattern of pPAR polarity that appeared upon reanimation. Specifically, initially bipolar embryos restored PAR-2 localization to both poles, while monopolar embryos restored PAR-2 localization to the pole at which it was initially enriched (Figure 5C, 5D). Thus, the distribution of persistent aPAR clusters in anoxia accurately predicts the pattern of polarity re-establishment.

Finally, we asked whether recovery of polarity after anoxia was dependent on persistent asymmetry of aPAR clusters. To test this, we compared the ability of embryos expressing oligomeric and monomeric PAR-3 variants to restore polarity upon reanimation. Polarity recovery was assessed by asymmetry of the first division, reflected in differences in daughter cell size (AB vs. P1) and cell-cycle timing, with AB normally dividing before P1 (Figure 5E, compare RitC with WT) (Delattre and Goehring, 2021; Rose and Gonczy, 2014).

For oligomeric PAR-3 variants, PAR-3(WT) and PAR-3(2mer), which remain asymmetric during anoxia (see Figure 4A), both size asymmetry and division asynchrony were normal following reanimation, consistent with accurate re-establishment of polarity (Figure 5E-G). By contrast, for PAR-3(RitC), which is polarized normally in normoxia but becomes symmetric in anoxia (see Figure 4A), both size asymmetry and division asynchrony were defective (Figure 5E-G). These defects were comparable to those seen for PAR-3(Δ69-82), which fails to become asymmetric due to defective membrane association (note *par-3(Δ69-82)* embryos already show significant defects in normoxia).

To confirm that defects in *par-3(RitC)* embryos were associated with a failure to properly restore polarity after anoxia, we examined the localization of PAR-2 and PAR-6. Unlike WT, in *par-3(RitC)* embryos the distribution of PAR-2 upon reanimation was highly variable and did not reflect the initially normal PAR-2 distribution at the onset of anoxia (Figure 5H,I, WT vs RitC). PAR-2 was typically enriched in lateral bands (21/24), often failed to coalesce into a well-defined posterior domain (15/24), and in 10/24 embryos was enriched in bipolar caps (Figure 5H, Figure S5F).

Taken together, these data lead us to conclude that the consolidation of PAR-3 and other aPARs into a stable and asymmetric population of membrane-associated clusters is required to encode a persistent memory of polarity during suspended animation. The resulting persistent memory state poises embryos to rapidly re-establish PAR polarity, undergo timely asymmetric division, and resume their normal developmental trajectory.

## Discussion

Current models for PAR polarity are generally formulated as information-activity feedback circuits, in which PAR proteins provide spatial cues to modulate local patterns of kinase and GTPase activity. These patterns of activity in turn shape the distribution of PAR proteins within the cell. However, while such models can explain the role of protein dynamics in the establishment and maintenance of polarity under normal conditions, as active biochemical systems, such networks are sensitive to perturbations such as anoxia, in which energy (and thus active processes) becomes limiting. Under these conditions, loss of activity would be expected to drive a catastrophic loss of information, i.e. polarity loss. Indeed, prior work has shown polarity markers are often disrupted during energy-compromised states (Dong et al., 2015; Lu et al., 2022) and as we show here pPARs depolarize in anoxia. However, our data suggest that the PAR polarity network is in fact able to retain critical spatial information in anoxia by transitioning into a physically encoded memory state. This state allows cells to preserve spatial information under conditions in which it cannot be actively maintained by energy-dependent network dynamics and then use this information to rapidly re-establish polarity once energy is returned to the system (Figure 6).

**Figure 6:**
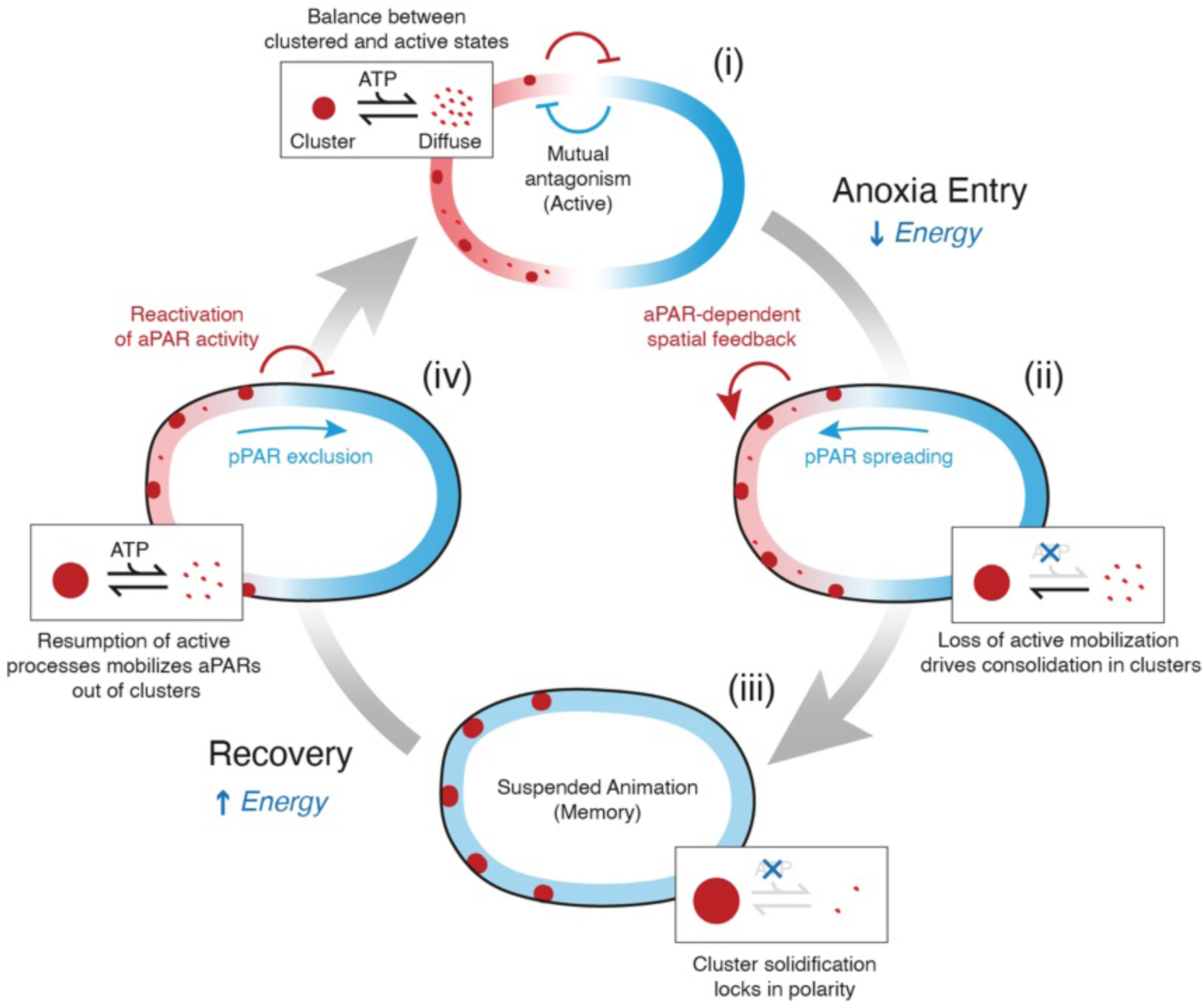
A model for maintaining polarity information during anoxia: (i) Mutual exclusive PAR domains are maintained by active mechanisms including mutual antagonism between PAR proteins. Here aPAR asymmetry is actively maintained by the posterior PAR proteins. PAR-3 dynamically shifts between clustered and diffuse states. (ii) Withdrawal of energy leads to loss of mutual antagonism. Consequently, pPARs are able to invade anterior but are unable to displace aPARs, which consolidate within anterior clusters in a manner that is biased by aPAR-dependent feedback. (iii) In long-term anoxia, aPARs, driven by PAR-3, are immobilized within solid-like clusters to lock-in a ‘memory’ of polarity. (iv) Re-animation mobilizes aPARs from PAR-3 clusters, which, coupled to re-activation of mutual antagonism, allows re-establishment of normal, mutually exclusive PAR domains.

Mechanistically, this memory state is linked to dynamics of oligomerization-dependent PAR-3 clusters. Initially, PAR-3 clusters are restricted to the anterior through active cortical actomyosin flow and biochemical input from the rest of the PAR network. Energy is also expended to destabilize clusters, at least in part via phosphorylation of PAR-3 by PLK-1, thereby maintaining PAR-3 in a dynamic responsive state. However, as oxygen becomes limiting and ATP levels drop, PAR-3 clusters become progressively stabilized. Ultimately, with extended energy depletion, dynamic exchange of PAR-3 into and out of clusters ceases in a process that resembles solidification, which together with the reduced mobility of clusters effectively locks in the spatial distribution of PAR-3 to create a long-term energy-independent polarized state. Upon reanimation, increasing ATP restores the activity of PAR proteins and their flux out of clusters to regenerate dynamic and responsive polarized activity patterns. Whether this behavior is linked to the reported ability of PAR-3 to phase separate in other systems is not clear, but at minimum suggests that PAR-3 has the capacity to adopt solid or gel-like behavior (Benton and Johnston, 2003; Kono et al., 2019; Liu et al., 2020).

While this work has focussed on the role of PAR-3 clusters as a potential long-term spatial memory during suspended animation, the same properties of clusters that enable memory during anoxia play important roles under normal conditions. As we and others have argued, persistent features such as long-lived protein clusters or oligomers allow cells such as the *C. elegans* zygote to integrate the effects of asymmetric transport over extended time periods, enhancing transport-dependent segregation, while also rendering the resulting asymmetries more persistent (Illukkumbura et al., 2023; Lang and Munro, 2022). Such time integration may also buffer spatiotemporal fluctuations in polarizing signals or the activity of the kinases and GTPases. More broadly, persistent cytoskeletal assemblies, including septins and intermediate filaments like vimentin and keratins, as well as extracellular matrix organization have been implicated in the persistence of polarity and spatial memory in mammalian cells and tissues. Taken together, these observations suggest that stable protein assemblies may represent a general mechanism by which polarity networks encode persistent spatial memory (Boubakar et al., 2017; Gan et al., 2016; Lim et al., 2020; Van Haastert, 2021). Such assemblies may function analogously to the metastable “ghost” states that have been proposed for signaling networks controlling cell migration that are thought to balance persistence and adaptability in the context of chemoattractant-guided cell migration (Nandan et al., 2022).

In conclusion, we suggest that regulation of PAR-3 clustering dynamics allows cells to balance responsiveness and stability within cell polarity circuits. By modulating the stability of PAR-3 clusters, cells can optimize the persistence of polarity in response to changing physiological demands, of which suspended animation represents an extreme case.

## Methods

### *C. elegans* strains and culture conditions

*C. elegans* were grown on nematode growth media (NGM) plates, seeded with OP50 bacterial lawns. All lines were kept at 20℃, unless stated otherwise. All strains used in this study are listed in table S1.

### Generating CRISPR strains

CRISPR-Cas9 mutations were introduced following the protocol previously published by (Arribere et al., 2014). Briefly, tracrRNA (IDT DNA, 0.5 μL at 100 μM) and crRNA(s) for the target (IDT DNA, 2.7 μL at 100μM) were incubated with duplex buffer (IDT DNA, 2.8μL) for 5 min at 95°C. Before injection, Cas9 (IDT DNA, 0.5μL at 10mg/mL) and a co-CRISPR injection marker (either a dpy-10 or unc-58) were incubated with the crRNA/tracrRNA mix for 15 min at 37°C. Mixture was then centrifuged for 10 min (13,000 rpm) before injection in young gravid adults. Mutants were then identified by PCR and the sequence confirmed by sequencing.

### Dissection and mounting of embryos in normoxia conditions

Worms were dissected on a coverslip (22×22 mm) in a 10-12 µl drop of Egg buffer (EB) (118 mM NaCl, 48 mM KCl, 2 mM CaCl2 2 mM MgCl2, 25 mM HEPES, pH 7.3), containing either 18.8 µm or 20 µm polystyrene beads (Polysciences, Inc), for cortex or midplane imaging, respectively. Embryos were then mounted between the coverslip and a large coverslip (1.5H, 60 x 24 mm, high-precision), and sealed with VALAP (1:1:1, vaseline:lanolin:paraffin wax).

### Anoxia/recovery experiments

For anoxia experiments, unless stated otherwise, embryos were incubated in a solution of PCA (2.5 mM, Protocatechuic acid/3,4-Dihydroxybenzoic acid, Sigma-Aldrich) and PCD (100 nM, Protocatechuate 3,4-Dioxygenase, Sigma-Aldrich) in EB. PCA was prepared as a stock solution of 100 mM, by dissolving PCA powder in distilled water, and adding NaOH (1M) slowly until complete dissolution of PCA was observed. The PCA solution was maintained at 4℃ and freshly prepared every week. A stock solution of PCD (5 µM) was maintained at - 20℃ and prepared in a buffer of 50% glycerol, 50 mM KCl, 1 mM EDTA, 100mM Tris-HCl (pH = 8). When mounting embryos for anoxia recovery experiments, parafilm was placed between the small coverslip and the VALAP on both sides of the coverslip. Depending on the experiment, embryos were left in anoxic conditions either overnight or for at least 2h in anoxia before recovery. For recovery, parafilm was then removed, disrupting the VALAP seal, and creating two open sides to allow exchange of buffer with fresh EB.

### Other energy depleting experiments (Sodium sulfite and anoxic bags)

For energy depleting experiments using sodium sulfite, embryos were mounted in 1M of sodium sulfite solution prepared with EB and polystyrene beads. Embryos were arrested for at least 2h, before recovery with fresh EB. For recovery, fresh EB was then placed on one side of the slide and exchanged via capillary action, as mentioned before.

For energy depletion using anoxic bags (Microbiology Anaerocult ® P, Merck Millipore), young adult worms were transferred to OP50 seeded NGM plates. These plates were then placed in sealed anoxic bags for at least 12h. The environment in these bags slowly becomes anoxic, which can be monitored by the indicator (Microbiology Anerotest ®, Merck Millipore), causing embryos to arrest and adult worms to immobilise. The following day, the worms were removed from the anoxic bags and quickly dissected in fresh EB to induce re-animation of the embryonic cell cycle.

### RNAi culture conditions

RNAi depletion was performed by feeding, as described previously (Kamath and Ahringer, 2003). Briefly HT115(DE3) bacterial feeding clones were inoculated into LB liquid cultures and grown overnight at 37°C. Culture was spiked with IPTG (final concentration 10 µM) before seeding into NGM plates (supplemented with 25µg/ml Carbinicillin and 1mM IPTG). After 24 h of incubation at 20°C, L3/L4 larvae were added to these RNAi feeding plates, for 24-48h, depending on the RNA being depleted. For the RNAi experiments in figure 1, chin-1 RNAi was performed for at least 45h. For complete PAR-2 depletion, at least 24h of RNAi was performed. For Figure 2, a range of 14-39h was used for par-3 RNAi, and a range of 19-39h was used for cdc-42 RNAi. For Figure 4, to achieve a complete RNAi depletion, at least 42h of cdc-42 RNAi were performed, and at least 46h of RNAi was performed for pkc-3 and par-6 RNAi. In Figure 5, for air-1 RNAi, at least 24h of RNAi was performed.

### Survival/hatching assay

For long term survival assays, embryos were recovered after anoxia and followed until larva stage. For this, embryos were mounted in a PCA/PCD anoxic buffer and left ON at 20°C (5-6 worms dissected) using a 18×18 mm and a 24×60 mm coverslip. The morning after, embryos were recovered as described above (parafilm is removed and fresh EB is washed in, several times, for at least 30-40 min). Using a scalpel, the VALAP seal was broken and the two coverslips were carefully separated, carefully adding more EB to the embryos, so these didn’t dry. Oocytes were manually removed using a needle, as these would not mature and would bias the rate of recovery. Embryos were then incubated ON in EB at 20°C (in a humid chamber). After 24h, the ratio of embryos (non-recovered) vs larvae (recovered) was quantified.

### Microscopy live imaging: midplane imaging

Midplane imaging was carried out in Nikon TiE, using a 100x/1.40 NA oil objective. The microscope was equipped with a custom X-Light V1 spinning disk system (CrestOptics, S.p.A.) with 50 μm slits, 488, 561 fiber-coupled diode lasers (Obis) and Photometrics Evolve 512 Delta EMCCD camera. To control the microscope, Metamorph was used (Molecular Devices), which was configured by Cairn Research. Filter sets were from Chroma (Bellows Falls, VT): ZT488/561rpc, ZET405/488/561/640X, ET535/50m, ET630/75m. Multi-dimensional acquisition was used to sequentially acquire images in the following channels: DIC, GFP (ex488, ET535/50m), RFP (ex561, ET630/75m). Autofluorescence, when recorded, was acquired with ex488, ET630/75m. All embryos were imaged at 20°C, using a temperature collar. For long anoxia periods, embryos were imaged every 5 min for the first 3h, and then every 30 min for a total time of 12-15h. For short anoxia periods, embryos were imaged every 1-2 min. Imaging in anoxia started as soon as possible after embryos were mounted (this corresponds to “t0”). Recovery was imaged for at least 2h in anoxia, or until all embryos divided at least twice, as described above. Embryos were imaged every 2 min during recovery.

### Microscopy live imaging: cortex imaging

Cortex imaging was carried out using a 100x 1.49 NA TIRF objective. Nikon TiE microscope equipped with an iLas2 TIRF unit (Roper) and a custom-made field stop, 488 or 561 fiber coupled diode lasers (Obis), and an Evolve 512 Delta EMCCD camera (Photometrics). Metamorph (Molecular Devices) was used to control the microscope (configured by Cairn Research). Filter sets were from Chroma: ZT488/561rpc, ZET488/561x, ZET488/561m, ET525/50m, ET630/75m, ET655LP. Images were captured in bright field, GFP/mNG (ex488/ZET488/561m), RFP/mKate/mCherry/Halo (ex561/ZET488/561m). For anoxia entry, embryos were imaged every 1 min. For single molecule tracking, frames were acquired every 33 ms, for a total of 1500 frames. For cluster tracking, frames were acquired every 1 s for 300 frames at 100 ms exposure. For colocalization analysis, a 1.5x optovar was used.

### FRAP - imaging conditions

To examine the mobility of the clusters by FRAP, anoxia was induced for the indicated period of time and embryos were then imaged (10% laser power, 100 ms) for 10 min before the FRAP event, and then followed for 50 min, imaging every 1 min. During the FRAP event, 5 clusters (or plasma membrane regions in the case of PAR-2) were simultaneously bleached, by using 5 squares of 10×10 with 100% laser power for 5 pulses.

### Image analysis - quantification of plasma membrane profile

Raw or autofluorescence-corrected (SAIBR, (Rodrigues et al., 2022)) images were used for quantification. N2 embryos were used for the autofluorescence calibration. Cortical concentrations were measured as published before (Gross et al., 2019; Ng et al., 2022; Reich et al., 2019). Briefly, a 100-pixel-wide (15.5 μm) line was traced following the membrane around the embryo, which was then computationally straightened. A 20-pixel-wide (3.1 μm) rolling average filter was then applied to the straightened plasma membrane. Local plasma membrane concentrations were calculated as the amplitude of the Gaussian component, at each position of the straightened line. For SAIBRed images, an error function component was also calculated, representing the cytoplasmic signal. For live imaging movies, a new ROI was traced for each frame. Only embryos in maintenance were analysed.

ASI was calculated using the equation: ASI = (A – P) / (2 * (A + P)). Anterior (A) and posterior (P) domains were found based on positional information, and are calculated as the average of the intensity of the plasma membrane profiles of the 30% anterior-most and posterior-most, respectively (Rodriguez et al., 2017). ASIs were normalised to the first frame: embryo is polarised when ASI = 1, and polarity loss corresponds to ASI = 0.

Plasma membrane profiles were calculated by averaging profiles from the upper and lower hemispheres of midplane embryo images. Two consecutive timepoints were averaged to reduce technical noise (for the start of anoxia: frame 1 and 2 were used, for the end of anoxia: last frame in anoxia and first frame in recovery were used). To facilitate comparisons, the plasma membrane intensity of all proteins was normalised to the first frame of anoxia. Plots and the smoothened conditional means were generated in R.

### Image analysis - DIC image correlation coefficient

Correlation coefficients during anoxia and recovery were calculated using the Image CorrelationJ Fiji plugin (Chinga and Syverud, 2007). Briefly, correlations were calculated from consecutive DIC image frames and the resulting values of R^2^ normalised to the last 3h in anoxia. Plots and mean curves were generated in R.

### Image analysis - cluster quantification

For Figure 2, 10 consecutive frames were acquired. Only clusters present in all frames were considered with the mean mass calculated across frames for each cluster. Total cluster intensity (sum of cluster masses) was normalised to the corresponding embryo area. For Figure 2 I, single frames were used. For the RNAi experiments in Figure 4, cluster number was normalised to the respective domain area (either anterior or posterior). For ASI calculations with TIRF data, the total cluster intensity of each domain (normalised to the respective domain area) was considered, and the following equation was used: (anterior - posterior) / (anterior + posterior). For cluster number ratio calculations, the ratio of clusters in posterior/anterior domain was found (each normalised to the respective domain area). All normalisations and plotting were performed in R.

### Image analysis - embryo recovery from anoxia

For embryo recovery from long term anoxia (> 12h) in Figure 1, embryos were recovered as mentioned above and monitored by live-imaging for at least 2 rounds of cell division (4-cell stage). Only 1-cell embryos were used for this quantification. Division was classified as “div.” (if divisions behave like normoxic WT development), “abnormal div.” (if cells don’t follow the WT development, AB and P1 divide synchronously for example) and “no div.” (if embryos don’t divide after recovery).

All plots were generated in R.

### Image analysis - AB/P1 cell division

AB/P1 cell division was classified as “asynchronous” when AB and P1 cells divide asynchronously, as expected in normal WT development (AB cell divides first than P1 cell). If AB and P1 cells divide simultaneously, AB/P1 was classified as “synchronous”. For each cell, the start of cell division was assessed by the start of furrow ingression. All plots were generated in R.

### Image analysis - P1 cell size

The P1 cell size was measured on a midplane cross-section, before either cell starts to round up (AB/P1 cell-cell contact is straight). For this, the area of AB and P1 blastomeres was found in Fiji using either manually traced ROIs or through semi-automated segmentation as in (Rodrigues et al., 2022; Rodrigues et al., 2024). P1 size was then calculated by dividing the P1 size by the area of the entire embryo. For Figure 5, a pairwise t test was performed in R, using the Bonferroni method to adjust p values for multiple comparisons and taking normoxia WT embryos as the reference group. All plots were generated in R.

### Image analysis - co-localisation (Pearson’s coefficient)

To quantify colocalization, dual channel images were captured preprocessed using a Median filter (1 px) and subject to rolling circle (50 px) background subtraction to suppress autofluorescence. Pre-processed red/green image pairs were then analyzed over an ROI encompassing the anterior domain using the BIOP JACOB plugin employing automatic thresholding (RenyiEntropy) and Costes randomization (2D, no thresholding, block size:5, shuffling iterations (100). (https://c4science.ch/w/bioimaging_and_optics_platform_biop/image-processing/imagej_tools/jacop_b/).

### Image analysis - Single Particle Detection and Tracking

The Trackpy package (https://github.com/soft-matter/trackpy) was used to carry out single molecule and cluster tracking using custom Python code as described (Hubatsch et al., 2019; Illukkumbura et al., 2023) (https://github.com/lhcgeneva/SPT). Briefly, particles are localised by fitting local intensity peaks (in individual frames) to a Gaussian point spread function, using the Crocker-Grier algorithm as described (Illukkumbura et al., 2023). For single particle tracking, the following parameters were used to optimize tracking errors: feature size = 5 pixels, memory = 0 frames, maximum distance features can move between frames = 3 pixels, minimum track length = 11 frames. For cluster tracking, the following parameters were used: maximum distance features can move between frames = 5 pixels, minimum track length = 5 frames. The same package was used to measure the cluster intensity during anoxia. Curves were fitted to a one-phase decay curve and plots were generated in Prism.

### Image analysis - FRAP

Embryos were aligned before image analysis to minimize microscope stage drift using the StackReg plugin in Fiji (Thevenaz et al., 1998). The intensity of the bleached areas was found using Fiji. For each embryo, the intensity of the background and 3 non-bleached clusters was taken as control for bleaching during imaging, respectively. The intensity of bleached clusters was normalized by the intensity of control clusters, with each corrected for background intensity in R. Pre-bleach values were normalized to 1. For each condition, the average FRAP curve and the fitted curve (one-phase association) was then calculated using Prism to obtain the timescale of recovery (tau) and the immobile fraction.

### Model

Simulations of polarity loss upon energy withdrawal were performed using a one-dimensional partial differential equation model, adapted from (Goehring et al., 2011b; Gross et al., 2019). Briefly, the model captures the membrane-bound diffusion, uniform recruitment and dissolution of anterior and posterior PARs assuming a well-mixed, finite cytoplasmic pool (*A* and *P*). aPARs and pPARs additionally undergo cross-repression by catalysing the membrane dissolution of one another (*k_AP_*, *k_PA_*), which model phosphorylation (energy) dependent processes. To model polarity loss into anoxia, we triggered polarisation via transient relief of aPAR-mediated pPAR removal (*k_PA_*), and solved the above system to steady-state, using a finite difference schema (normoxia). This then serves as an initial condition for anoxic simulations, where the cross-repression terms are set instantaneously to zero. We performed a parameter scan of concentration trajectories along the dimensionless parameter (*D_A_/(k_off,A_L^2^*), and used this to build the two-dimensional phase space in Figure 1B, C. Time to polarity loss is calculated as the time for asymmetry (ASI) to be reduced by 90%. Model construction, equations and simulation details are provided in the Supplementary Model Description.

---

**Table S1.**
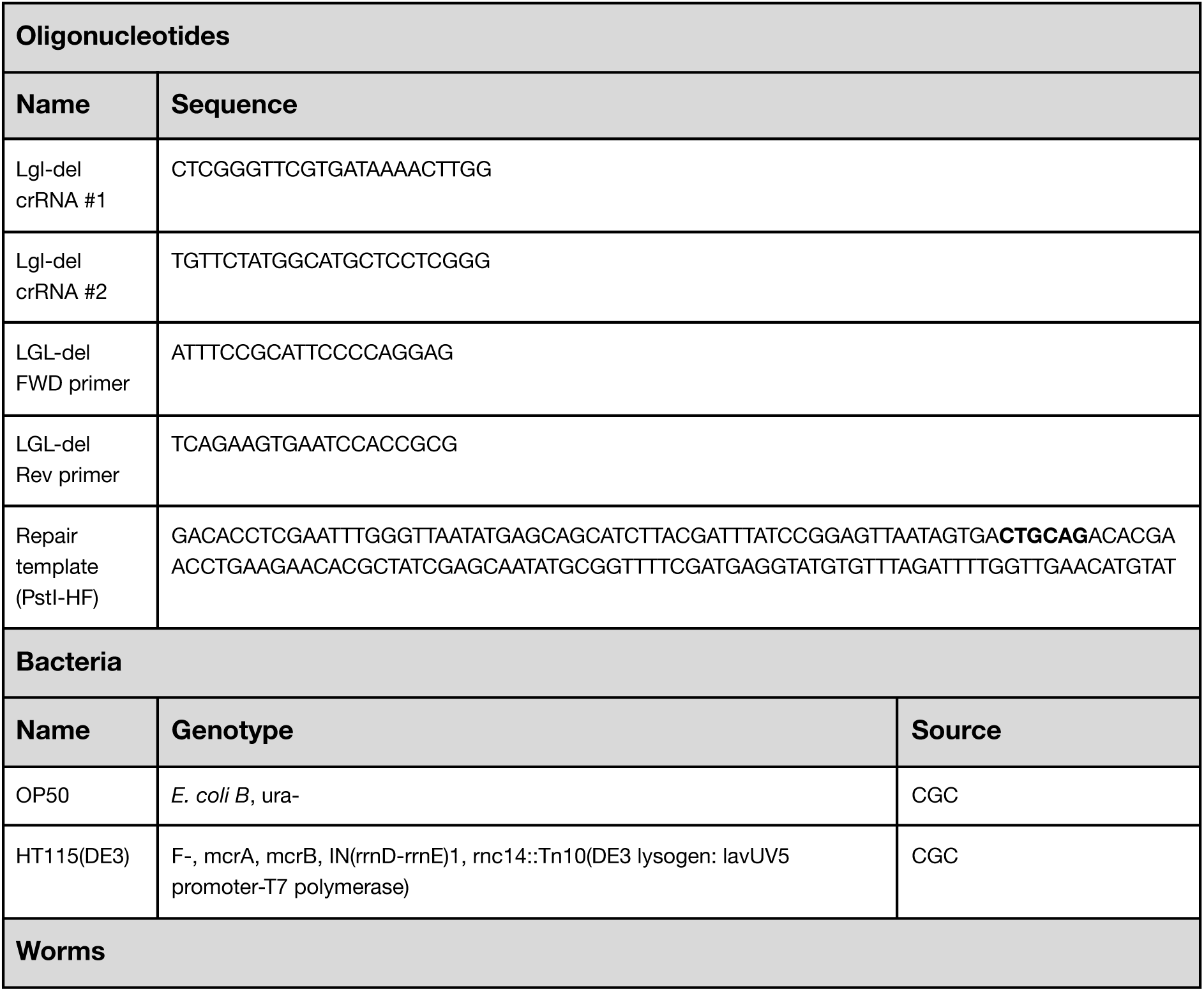
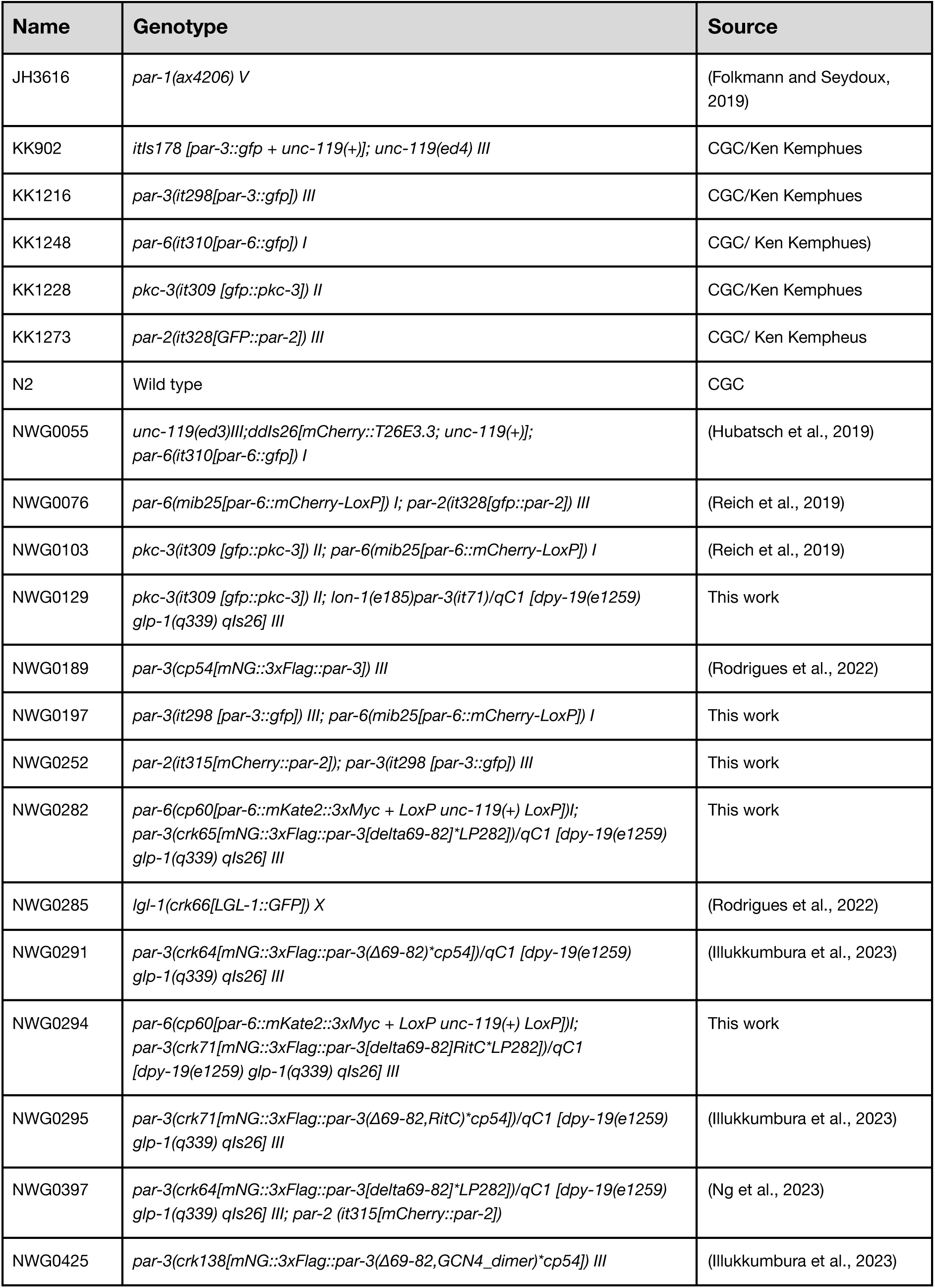
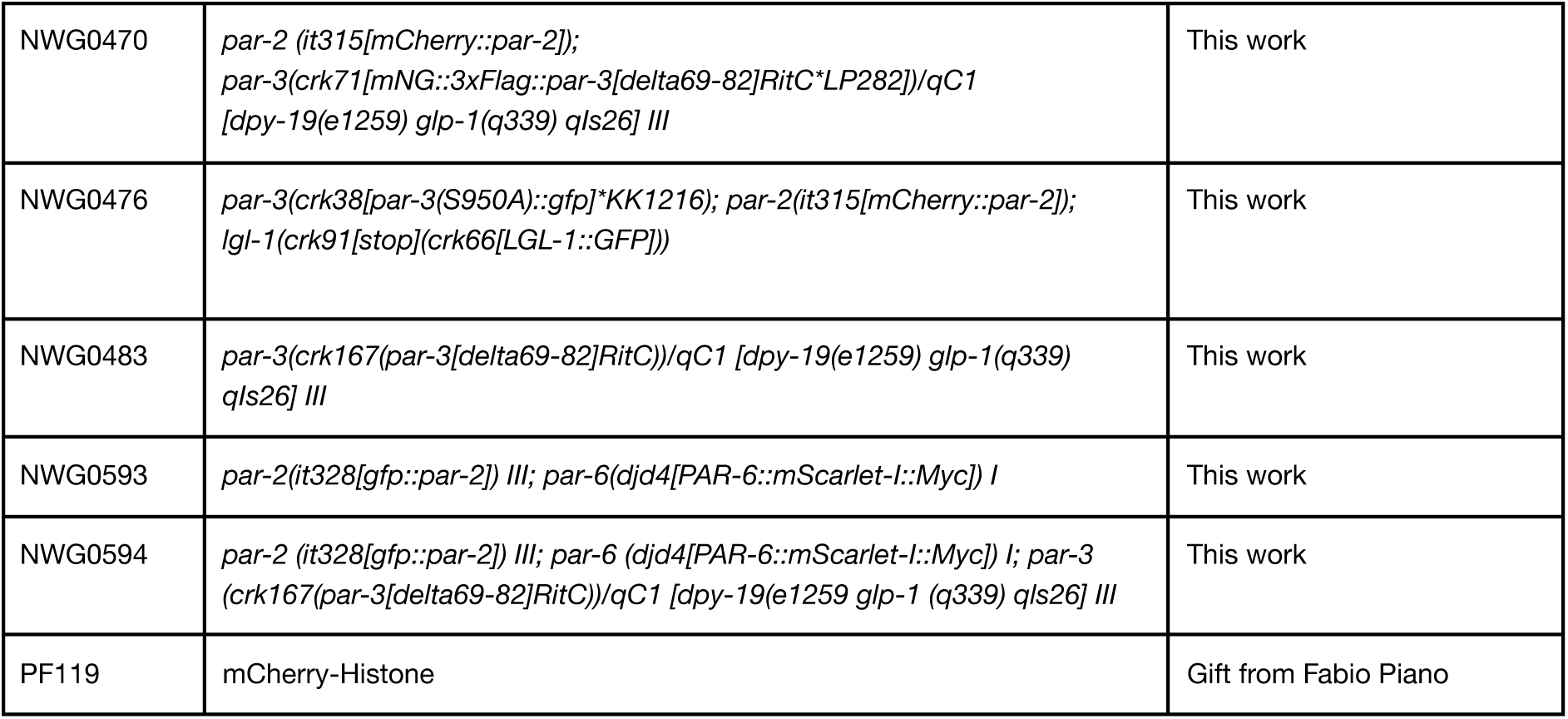
Resources and Reagents.

## Contributions

Conceptualization: J.P., N.W.G.; Methodology: J.P., N.W.G.; Formal analysis: J.P., J.C.S.; Investigation: J.P., N.H., J.C.S.; Resources: J.P., N.H.; Writing – original draft preparation: J.P., N.W.G.; Writing – review and editing: J.P., J.C.S., B.B., S.B., N.W.G.; Supervision: B.B., S.B., N.W.G.; Project administration: B.B., S.B., N.W.G.; Funding acquisition: B.B., S.B., N.W.G.

## Acknowledgements

We thank the Goehring Lab for comments on the manuscript. Strains were provided by the *Caenorhabditis* Genome Center (CGC), which is funded by NIH Office of Research Infrastructure Programs (P40 OD010440). This work was supported by the Francis Crick Institute, which receives its core funding from Cancer Research UK (CC1051, CC2119), the UK Medical Research Council (CC1051, CC2119), and the Wellcome Trust (CC1051, CC2119), a summer studentship from the Institute for Physics of Living Systems (UCL) and a Boehringer Ingelheim Fonds PhD fellowship (to J.C.S.), and a NIH R35 GM143042 (S.B.).

## Data Availability

Source code and documentation will be made available on GitHub/FigShare upon publication.

## Supplemental Materials

**Supplemental Text: Supplementary Model Description**

**Movie S1: PAR-2 and PAR-3 localization during transition into anoxia.** PAR-3::GFP and GFP::PAR-2 expressing embryos imaged via spinning disk confocal. Time(min) indicated relative to anoxia entry.

**Movie S2: Growth and solidification of PAR-3 clusters during transition into anoxia.** PAR-3::GFP expressing embryos imaged via TIRF. Time (min) relative to anoxia entry.

**Movie S3: Recruitment of PAR-6 to PAR-3 cluster during transition into anoxia.** PAR-3::GFP (red), PAR-6::mKate (cyan) expressing embryos imaged via TIRF. Time (min) relative to anoxia entry.

**Movie S4: PAR-3 cluster behavior during transition into anoxia (WT vs CDC-42).** PAR-3::GFP expressing embryos subject to *ctl* (Wild-type) or *cdc-42(RNAi)* imaged via TIRF. Time(min) relative to anoxia entry.

**Movie S5: Recovery of polarity after anoxia requires PAR-3 clustering.** Wild-type or *par-3(RitC)* embryos expressing PAR-6::mScarlet / GFP::PAR-2 shown during anoxia entry and subsequent recovery upon return to normoxia imaged via spinning disk confocal. Time(min) relative to anoxia entry, return to normoxia shown.

## References

Aceto, D., Beers, M. and Kemphues, K. J. (2006). Interaction of PAR-6 with CDC-42 is required for maintenance but not establishment of PAR asymmetry in C. elegans. Dev. Biol. 299, 386–397.

Aitken, C. E., Marshall, R. A. and Puglisi, J. D. (2008). An Oxygen Scavenging System for Improvement of Dye Stability in Single-Molecule Fluorescence Experiments. Biophys. J. 94, 1826–1835.

Anderson, D. C., Gill, J. S., Cinalli, R. M. and Nance, J. (2008). Polarization of the C. elegans Embryo by RhoGAP-Mediated Exclusion of PAR-6 from Cell Contacts. 320, 1771–1774.

Arribere, J. A., Bell, R. T., Fu, B. X. H., Artiles, K. L., Hartman, P. S. and Fire, A. Z. (2014). Efficient Marker-Free Recovery of Custom Genetic Modifications with CRISPR/Cas9 in Caenorhabditis elegans. Genetics 198, 837–846.

Beatty, A., Morton, D. G. D. and Kemphues, K. (2013). PAR-2, LGL-1 and the CDC-42 GAP CHIN-1 act in distinct pathways to maintain polarity in the C. elegans embryo. Dev. Camb. Engl. 140, 2005–2014.

Benton, R. and Johnston, D. S. (2003). A Conserved Oligomerization Domain in Drosophila Bazooka/PAR-3 Is Important for Apical Localization and Epithelial Polarity. Curr. Biol. 13, 1330–1334.

Benton, R. and St Johnston, D. (2003). Drosophila PAR-1 and 14-3-3 inhibit Bazooka/PAR-3 to establish complementary cortical domains in polarized cells. Cell 115, 691–704.

Betschinger, J., Mechtler, K. and Knoblich, J. A. (2003). The Par complex directs asymmetric cell division by phosphorylating the cytoskeletal protein Lgl. Nature 422, 326–330.

Bickler, P. E. and Buck, L. T. (2007). Hypoxia Tolerance in Reptiles, Amphibians, and Fishes: Life with Variable Oxygen Availability. Annu. Rev. Physiol. 69, 145–170.

Boothby, T. C., Tapia, H., Brozena, A. H., Piszkiewicz, S., Smith, A. E., Giovannini, I., Rebecchi, L., Pielak, G. J., Koshland, D. and Goldstein, B. (2017). Tardigrades use intrinsically disordered proteins to survive desiccation. Mol. Cell 65, 975–984.e5.

Boubakar, L., Falk, J., Ducuing, H., Thoinet, K., Reynaud, F., Derrington, E. and Castellani, V. (2017). Molecular Memory of Morphologies by Septins during Neuron Generation Allows Early Polarity Inheritance. Neuron 95, 834–851.e5.

Boyd, L., Guo, S., Levitan, D., Stinchcomb, D. T. and Kemphues, K. J. (1996). PAR-2 is asymmetrically distributed and promotes association of P granules and PAR-1 with the cortex in C. elegans embryos. Dev. Camb. Engl. 122, 3075–3084.

Buck, L. T. and Hochachka, P. W. (1993). Anoxic suppression of Na(+)-K(+)-ATPase and constant membrane potential in hepatocytes: support for channel arrest. Am. J. Physiol. 265, R1020–1025.

Calvi, I., Schwager, F. and Gotta, M. (2022). PP1 phosphatases control PAR-2 localization and polarity establishment in *C. elegans* embryos. J. Cell Biol. 221, e202201048.

Campanale, J. P., Sun, T. Y. and Montell, D. J. (2017). Development and dynamics of cell polarity at a glance. J. Cell Sci. 130, 1201–1207.

Chang, Y. and Dickinson, D. J. (2022). A particle size threshold governs diffusion and segregation of PAR-3 during cell polarization. Cell Rep. 39, 110652.

Chinga, G. and Syverud, K. (2007). Quantification of paper mass distributions within local picking areas. Nord. Pulp Pap. Res. J. 22, 441–446.

Cowan, C. R. and Hyman, A. A. (2004). Centrosomes direct cell polarity independently of microtubule assembly in C. elegans embryos. Nature 431, 92–96.

Cross, M. C. and Hohenberg, P. C. (1993). Pattern formation outside of equilibrium. Rev. Mod. Phys. 65, 851–1112.

Delattre, M. and Goehring, N. W. (2021). Chapter Nine - The first steps in the life of a worm: Themes and variations in asymmetric division in C. elegans and other nematodes. In Current Topics in Developmental Biology (ed. Jarriault, S.) and Podbilewicz, B.), pp. 269–308. Academic Press.

Dickinson, D. J., Schwager, F., Pintard, L., Gotta, M. and Goldstein, B. (2017). A Single-Cell Biochemistry Approach Reveals PAR Complex Dynamics during Cell Polarization. Dev. Cell 42, 416–434.e11.

Dong, W., Zhang, X., Liu, W., Chen, Y., Huang, J., Austin, E., Celotto, A. M., Jiang, W. Z., Palladino, M. J., Jiang, Y., et al. (2015). A conserved polybasic domain mediates plasma membrane targeting of Lgl and its regulation by hypoxia. J. Cell Biol. 211, 273–286.

Etemad-Moghadam, B., Guo, S. and Kemphues, K. J. (1995). Asymmetrically distributed PAR-3 protein contributes to cell polarity and spindle alignment in early C. elegans embryos. Cell 83, 743–752.

Fenelon, J. C., Banerjee, A. and Murphy, B. D. (2014). Embryonic diapause: development on hold. Int. J. Dev. Biol. 58, 163–174.

Foe, V. E. and Alberts, B. M. (1985). Reversible chromosome condensation induced in Drosophila embryos by anoxia: visualization of interphase nuclear organization. J. Cell Biol. 100, 1623–1636.

Folkmann, A. W. and Seydoux, G. (2019). Spatial regulation of the polarity kinase PAR-1 by parallel inhibitory mechanisms. Development 146, dev171116–dev171116.

Franzmann, T. M. and Alberti, S. (2019). Protein phase separation as a stress survival strategy. Cold Spring Harb. Perspect. Biol. 11, a034058.

Gan, Z., Ding, L., Burckhardt, C. J., Lowery, J., Zaritsky, A., Sitterley, K., Mota, A., Costigliola, N., Starker, C. G., Voytas, D. F., et al. (2016). Vimentin Intermediate Filaments Template Microtubule Networks to Enhance Persistence in Cell Polarity and Directed Migration. Cell Syst. 3, 252–263.e8.

Goehring, N. W. (2014). PAR polarity: From complexity to design principles. Exp. Cell Res. 328, 258–266.

Goehring, N. W., Hoege, C., Grill, S. W. and Hyman, A. A. (2011a). PAR proteins diffuse freely across the anterior–posterior boundary in polarized C. elegans embryos. J. Cell Biol. 193, 583–594.

Goehring, N. W., Trong, P. K., Bois, J. S., Chowdhury, D., Nicola, E. M., Hyman, A. A. and Grill, S. W. (2011b). Polarization of PAR Proteins by Advective Triggering of a Pattern-Forming System. Science 334, 1137–1141.

Goldbeter, A. (2018). Dissipative structures in biological systems: bistability, oscillations, spatial patterns and waves. Philos. Trans. R. Soc. Math. Phys. Eng. Sci. 376, 20170376.

Goldstein, B. and Macara, I. G. (2007). The PAR proteins: fundamental players in animal cell polarization. Dev. Cell 13, 609–622.

Gotta, M., Abraham, M. C. and Ahringer, J. (2001). CDC-42 controls early cell polarity and spindle orientation in C. elegans. Curr. Biol. 11, 482–488.

Gross, P., Kumar, K. V., Goehring, N. W., Bois, J. S., Hoege, C., Jülicher, F. and Grill, S. W. (2019). Guiding self-organized pattern formation in cell polarity establishment. Nat. Phys. 15, 293–300.

Hajeri, V. a, Trejo, J. and Padilla, P. a (2005). Characterization of sub-nuclear changes in Caenorhabditis elegans embryos exposed to brief, intermediate and long-term anoxia to analyze anoxia-induced cell cycle arrest. BMC Cell Biol. 6, 47–47.

Hannaford, M., Loyer, N., Tonelli, F., Zoltner, M. and Januschke, J. (2019). A chemical-genetics approach to study the role of atypical protein kinase C in Drosophila. Dev. Camb.

Hapke, R. Y. and Haake, S. M. (2020). Hypoxia-induced epithelial to mesenchymal transition in cancer. Cancer Lett. 487, 10–20.

Heimlicher, M. B., Bächler, M., Liu, M., Ibeneche-Nnewihe, C., Florin, E.-L., Hoenger, A. and Brunner, D. (2019). Reversible solidification of fission yeast cytoplasm after prolonged nutrient starvation. J. Cell Sci. 132, jcs231688.

Helena-Bueno, K., Chan, L. I. and Melnikov, S. V. (2024). Rippling life on a dormant planet: hibernation of ribosomes, RNA polymerases, and other essential enzymes. Front. Microbiol. 15, 1386179.

Hoege, C., Constantinescu, A.-T., Schwager, A., Goehring, N. W., Kumar, P. and Hyman, A. A. (2010). LGL Can Partition the Cortex of One-Cell Caenorhabditis elegans Embryos into Two Domains. Curr. Biol. 20, 1296–1303.

Hsu, S.-P. and Dickinson, D. J. (2026). Multivalent assembly of PAR-3/aPKC complexes establishes cell polarity in *Caenorhabditis elegans* zygotes. Proc. Natl. Acad. Sci. 123, e2509713123.

Hubatsch, L., Peglion, F., Reich, J. D., Rodrigues, N. T. L., Hirani, N., Illukkumbura, R. and Goehring, N. W. (2019). A cell-size threshold limits cell polarity and asymmetric division potential. Nat. Phys. 15, 1078–1085.

Hung, T. J. and Kemphues, K. J. (1999). PAR-6 is a conserved PDZ domain-containing protein that colocalizes with PAR-3 in Caenorhabditis elegans embryos. Dev. Camb. Engl. 126, 127–135.

Hurov, J. B., Watkins, J. L. and Piwnica-Worms, H. (2004). Atypical PKC phosphorylates PAR-1 kinases to regulate localization and activity. Curr. Biol. 14, 736–741.

Illukkumbura, R., Hirani, N., Borrego-Pinto, J., Bland, T., Ng, K., Hubatsch, L., McQuade, J., Endres, R. G. and Goehring, N. W. (2023). Design principles for selective polarization of PAR proteins by cortical flows. J. Cell Biol. 222, e202209111.

Joyner, R. P., Tang, J. H., Helenius, J., Dultz, E., Brune, C., Holt, L. J., Huet, S., Müller, D. J. and Weis, K. (2016). A glucose-starvation response regulates the diffusion of macromolecules. eLife 5, e09376.

Kamath, R. S. and Ahringer, J. (2003). Genome-wide RNAi screening in Caenorhabditis elegans. Methods San Diego Calif 30, 313–321.

Kapoor, S. and Kotak, S. (2019). Centrosome Aurora A regulates RhoGEF ECT-2 localisation and ensures a single PAR-2 polarity axis in C. elegans embryos. Development 146, dev174565–dev174565.

Khuc Trong, P., Nicola, E. M., Goehring, N. W., Kumar, K. V. V. and Grill, S. W. (2014). Parameter-space topology of models for cell polarity. New J. Phys. 16, 065009–065009.

Kirschner, M., Gerhart, J. and Mitchison, T. (2000). Molecular “vitalism.” Cell 100, 79–88.

Klinkert, K., Levernier, N., Gross, P., Gentili, C., von Tobel, L., Pierron, M., Busso, C., Herrman, S., Grill, S. W., Kruse, K., et al. (2019). Aurora A depletion reveals centrosome-independent polarization mechanism in Caenorhabditis elegans. eLife 8, 388918–388918.

Kono, K., Yoshiura, S., Fujita, I., Okada, Y., Shitamukai, A., Shibata, T. and Matsuzaki, F. (2019). Reconstruction of Par-dependent polarity in apolar cells reveals a dynamic process of cortical polarization. eLife 8, e45559–e45559.

Korotkevich, E., Niwayama, R., Courtois, A., Friese, S., Berger, N., Buchholz, F. and Hiiragi, T. (2017). The Apical Domain Is Required and Sufficient for the First Lineage Segregation in the Mouse Embryo. Dev. Cell 40, 235–247.e7.

Kumfer, K. T., Cook, S. J., Squirrell, J. M., Eliceiri, K. W., Peel, N., O’Connell, K. F. and White, J. G. (2010). CGEF-1 and CHIN-1 Regulate CDC-42 Activity during Asymmetric Division in the Caenorhabditis elegans Embryo. Mol. Biol. Cell 21, 266–277.

Lang, C. F. and Munro, E. (2017). The PAR proteins: from molecular circuits to dynamic self-stabilizing cell polarity. Dev. Camb. Engl. 144, 3405–3416.

Lang, C. F. and Munro, E. M. (2022). Oligomerization of peripheral membrane proteins provides tunable control of cell surface polarity. Biophys. J. 121, 4543–4559.

Lang, C. F., Maxian, O., Anneken, A. and Munro, E. (2026). Oligomerization and positive feedback on membrane binding stabilize PAR-3 asymmetries in the C. elegans zygote. Curr. Biol. 36, 1509–1524.e5.

Li, B., Kim, H., Beers, M. and Kemphues, K. (2010). Different domains of C. elegans PAR-3 are required at different times in development. Dev. Biol. 344, 745–757.

Lim, H. Y. G., Alvarez, Y. D., Gasnier, M., Wang, Y., Tetlak, P., Bissiere, S., Wang, H., Biro, M. and Plachta, N. (2020). Keratins are asymmetrically inherited fate determinants in the mammalian embryo. Nature 585, 404–409.

Liu, Z., Yang, Y., Gu, A., Xu, J., Mao, Y., Lu, H., Hu, W., Lei, Q. Y., Li, Z., Zhang, M., et al. (2020). Par complex cluster formation mediated by phase separation. Nat. Commun. 11,.

Lu, J., Dong, W., Tao, Y. and Hong, Y. (2021). Electrostatic plasma membrane targeting contributes to Dlg function in cell polarity and tumorigenesis. Development 148, dev196956.

Lu, J., Dong, W., Hammond, G. R. and Hong, Y. (2022). Hypoxia controls plasma membrane targeting of polarity proteins by dynamic turnover of PI4P and PI(4,5)P2. eLife 11, e79582.

Lynch, E. M., Kollman, J. M. and Webb, B. A. (2020). Filament formation by metabolic enzymes—a new twist on regulation. Curr. Opin. Cell Biol. 66, 28–33.

Maire, T., Allertz, T., Betjes, M. A. and Youk, H. (2020). Dormancy-to-death transition in yeast spores occurs due to gradual loss of gene-expressing ability. Mol. Syst. Biol. 16, e9245.

Maity, S. and Moschou, P. N. (2026). Biomolecular condensates as cellular memory modules: thermodynamic principles and plant stress adaptation. Biophys. J. 125, 12–28.

Moreira, S., Osswald, M., Ventura, G., Gonçalves, M., Sunkel, C. E. and Morais-de-Sá, E. (2019). PP1-Mediated Dephosphorylation of Lgl Controls Apical-basal Polarity. Cell Rep. 26, 293–301.e7.

Motegi, F. and Seydoux, G. (2013). The PAR network: redundancy and robustness in a symmetry-breaking system. Philos. Trans. R. Soc. Lond. B. Biol. Sci. 368, 20130010–20130010.

Motegi, F., Zonies, S., Hao, Y., Cuenca, A. A., Griffin, E. E. and Seydoux, G. (2011). Microtubules induce self-organization of polarized PAR domains in Caenorhabditis elegans zygotes. Nat. Cell Biol. 13, 1361–1367.

Munder, M. C., Midtvedt, D., Franzmann, T., Nüske, E., Otto, O., Herbig, M., Ulbricht, E., Müller, P., Taubenberger, A., Maharana, S., et al. (2016). A pH-driven transition of the cytoplasm from a fluid- to a solid-like state promotes entry into dormancy. eLife 5,.

Munro, E., Nance, J. and Priess, J. R. (2004). Cortical flows powered by asymmetrical contraction transport PAR proteins to establish and maintain anterior-posterior polarity in the early C. elegans embryo. Dev. Cell 7, 413–424.

Nance, J., Munro, E. M. and Priess, J. R. (2003). C. elegans PAR-3 and PAR-6 are required for apicobasal asymmetries associated with cell adhesion and gastrulation. Dev. Camb. Engl. 130, 5339–5350.

Nandan, A., Das, A., Lott, R. and Koseska, A. (2022). Cells use molecular working memory to navigate in changing chemoattractant fields. eLife 11, e76825.

Ng, K., Bland, T., Hirani, N. and Goehring, N. W. (2022). An analog sensitive allele permits rapid and reversible chemical inhibition of PKC-3 activity in C. elegans. MicroPublication Biol.

Ng, K., Hirani, N., Bland, T., Borrego-Pinto, J., Wagner, S., Kreysing, M. and Goehring, N. W. (2023). Cleavage furrow-directed cortical flows bias PAR polarization pathways to link cell polarity to cell division. Curr. Biol. 33, 4298–4311.e6.

O’Connell, K. F., Maxwell, K. N. and White, J. G. (2000). The spd-2 gene is required for polarization of the anteroposterior axis and formation of the sperm asters in the Caenorhabditis elegans zygote. Dev. Biol. 222, 55–70.

Padilla, P. A. and Ladage, M. L. (2012). Suspended animation, diapause and quiescence: arresting the cell cycle in C. elegans. Cell Cycle Georget. Tex 11, 1672–9.

Padilla, P. A., Nystul, T. G., Zager, R. A., Johnson, A. C. M. and Roth, M. B. (2002). Dephosphorylation of Cell Cycle–regulated Proteins Correlates with Anoxia-induced Suspended Animation in *Caenorhabditis elegans*. Mol. Biol. Cell 13, 1473–1483.

Parry, B. R., Surovtsev, I. V., Cabeen, M. T., O’Hern, C. S., Dufresne, E. R. and Jacobs-Wagner, C. (2014). The Bacterial Cytoplasm Has Glass-like Properties and Is Fluidized by Metabolic Activity. Cell 156, 183–194.

Prigogine, I. (1978). Time, Structure, and Fluctuations. Science 201, 777–785.

Protter, D. S. W. and Parker, R. (2016). Principles and properties of stress granules. Trends Cell Biol. 26, 668–679.

Reich, J. D., Hubatsch, L., Illukkumbura, R., Peglion, F., Bland, T., Hirani, N. and Goehring, N. W. (2019). Regulated Activation of the PAR Polarity Network Ensures a Timely and Specific Response to Spatial Cues. Curr. Biol. 29, 1911–1923.e5.

Robin, F. B., McFadden, W. M., Yao, B. and Munro, E. M. (2014). Single-molecule analysis of cell surface dynamics in Caenorhabditis elegans embryos. Nat. Methods 11, 677–682.

Rodrigues, N. T. L., Bland, T., Borrego-Pinto, J., Ng, K., Hirani, N., Gu, Y., Foo, S. and Goehring, N. W. (2022). *SAIBR* : A simple, platform-independent method for spectral autofluorescence correction. Development dev.200545.

Rodrigues, N. T. L., Bland, T., Ng, K., Hirani, N. and Goehring, N. W. (2024). Quantitative perturbation-phenotype maps reveal nonlinear responses underlying robustness of PAR-dependent asymmetric cell division. PLOS Biol. 22, e3002437.

Rodriguez, J., Peglion, F., Martin, J., Hubatsch, L., Reich, J., Hirani, N., Gubieda, A. G., Roffey, J., Fernandes, A. R., St Johnston, D., et al. (2017). aPKC Cycles between Functionally Distinct PAR Protein Assemblies to Drive Cell Polarity. Dev. Cell 42, 400–415.e9.

Rose, L. and Gonczy, P. (2014). Polarity establishment, asymmetric division and segregation of fate determinants in early C. elegans embryos. WormBook 1–43.

Rosswag De Souza, S., Böke, E. and Zaffagnini, G. (2025). Proteostasis in cellular dormancy: lessons from yeast to oocytes. Trends Biochem. Sci. 50, 646–662.

Sailer, A., Anneken, A., Li, Y., Lee, S. and Munro, E. (2015). Dynamic Opposition of Clustered Proteins Stabilizes Cortical Polarity in the C. elegans Zygote. Dev. Cell 35, 131–142.

Schisa, J. A. (2014). Effects of stress and aging on ribonucleoprotein assembly and function in the germ line. Wiley Interdiscip. Rev. RNA 5, 231–246.

Schrödinger, E. (1944). What is life? The physical aspect of the living cell. Cambridge,: Cambridge University Press.

Schumacher, J. M., Ashcroft, N., Donovan, P. J. and Golden, A. (1998). A highly conserved centrosomal kinase, AIR-1, is required for accurate cell cycle progression and segregation of developmental factors in Caenorhabditis elegans embryos. Dev. Camb. Engl. 125, 4391–402.

Stolpner, N. J., Manzi, N. I., Su, T. and Dickinson, D. J. (2023). Apical PAR protein caps orient the mitotic spindle in C. elegans early embryos. Curr. Biol. 33, 4312–4329.e6.

Tabuse, Y., Izumi, Y., Piano, F., Kemphues, K. J., Miwa, J. and Ohno, S. (1998). Atypical protein kinase C cooperates with PAR-3 to establish embryonic polarity in Caenorhabditis elegans. Dev. Camb. Engl. 125, 3607–3614.

Tang, X., Qu, L., Wilfling, F., Beck, F., Ernst, O. P., Schulman, B. A., Baumeister, W. and Enenkel, C. (2026). Metabolically regulated proteasome supramolecular organization in situ. Cell 189, 1153–1169.e16.

Thevenaz, P., Ruttimann, U. E. and Unser, M. (1998). A pyramid approach to subpixel registration based on intensity. IEEE Trans. Image Process. 7, 27–41.

Traweger, A., Wiggin, G., Taylor, L., Tate, S. A., Metalnikov, P. and Pawson, T. (2008). Protein phosphatase 1 regulates the phosphorylation state of the polarity scaffold Par-3. Proc. Natl. Acad. Sci. U. S. A. 105, 10402–10407.

Van Haastert, P. J. M. (2021). Short- and long-term memory of moving amoeboid cells. PLOS ONE 16, e0246345.

Van Voorhies, W. A. and Ward, S. (2000). Broad oxygen tolerance in the nematode Caenorhabditis elegans. J. Exp. Biol. 203, 2467–2478.

Wallenfang, M. R. and Seydoux, G. (2000). Polarization of the anterior-posterior axis of C. elegans is a microtubule-directed process. Nature 408, 89–92.

Wang, S.-C., Low, T. Y. F., Nishimura, Y., Gole, L., Yu, W. and Motegi, F. (2017). Cortical forces and CDC-42 control clustering of PAR proteins for Caenorhabditis elegans embryonic polarization. Nat. Cell Biol. 19, 988–995.

Watts, J. L., Etemad-Moghadam, B., Guo, S., Boyd, L., Draper, B. W., Mello, C. C., Priess, J. R. and Kemphues, K. J. (1996). par-6, a gene involved in the establishment of asymmetry in early C. elegans embryos, mediates the asymmetric localization of PAR-3. Dev. Camb. Engl. 122, 3133–3140.

Zhao, P., Teng, X., Tantirimudalige, S. N., Nishikawa, M., Wohland, T., Toyama, Y. and Motegi, F. (2019). Aurora-A Breaks Symmetry in Contractile Actomyosin Networks Independently of Its Role in Centrosome Maturation. Dev. Cell 48, 631–645.e6.

